# Ground-truth *in silico* spike-ins reveal limits of microbiome biomarker recovery in colorectal cancer

**DOI:** 10.64898/2026.08.02.742082

**Authors:** André Salgado, Catarina Rosa Tomaz, Ana Teresa Freitas, Ana Santos Almeida

## Abstract

Microbiome-based biomarkers have been proposed for colorectal cancer (CRC), yet candidate taxa are often interpreted without knowing whether taxonomic profiling workflows can reliably detect and quantify them in human samples. Existing ground-truth studies commonly rely on simplified communities that do not preserve the biological and technical complexity of clinical stool metagenomes. We hypothesized that weak CRC-associated signals, particularly those relevant to early-stage disease, may be missed through analytical non-recovery rather than biological absence.

We developed an *in silico* spike-in framework that embeds CRC-associated signals into clinical stool metagenomes. Ten taxa were introduced individually at six fractions (0.01–5%) or as an equally weighted community at seven total fractions (0.01–10%; effective per-taxon fractions, 0.001–1%), generating 5,770 spike-in metagenomes from 310 samples. The resulting metagenomes were profiled with Kraken2/Bracken and MetaPhlAn 4 to quantify detection, abundance accuracy, false-positive signals, biomarker recovery, and calibration against a known ground truth.

Recovery depended strongly on workflow, taxon, abundance, and clinical background. At 0.01%, four taxa- *F. nucleatum*, *P. micra*, *P. stomatis*, and *P. intermedia*-showed good recovery in 85–90% of samples under Kraken2/Bracken, whereas none achieved good recovery in at least 50% under MetaPhlAn 4. Greater low-abundance recovery was accompanied by a broader artefact-prone background (54.9% versus 0.5% of non-target taxa). Artefact-prone taxa accounted for 96.3% and 100% of enriched off-target differential-abundance calls, respectively. Spike-in-derived artefact exclusion substantially reduced off-target detections where present, while abundance-response modelling provided proof-of-principle correction of systematic abundance distortions in both evaluated configurations.

Overall, the evaluated profiling configurations demonstrate that known low-abundance CRC-associated signals can be missed or distorted across a complete biomarker-discovery pipeline. Analytical non-recovery may cause early-detection biomarkers to be missed rather than indicate biological absence. Importantly, artefact-aware filtering and abundance calibration show that these limitations can be partially overcome. Improvements in taxonomic profiling may help bring reliable microbiome-based CRC diagnostics closer to clinical application.

## 1 Introduction

Microbiome-based biomarkers are a promising approach for non-invasive cancer detection, but their use in clinical practice is still limited by reproducibility. This challenge is particularly important in colorectal cancer (CRC). Several studies have reported associations between stool microbial composition and CRC status [1–6], but candidate taxa often show variable performance across cohorts, disease stages, and analytical workflows [7–9]. This problem is especially relevant for precancerous colorectal adenomas, where microbial changes are often small, sparse, and present in complex host-associated communities. Recent pooled analyses showed that microbiome-based classification of adenoma versus controls can be close to chance level, with AUC values close to 0.5 [5]. These results raise a practical question for CRC microbiome biomarker studies: when a reported disease-associated taxon is not detected, or does not contribute to classification, is this because the signal is biologically absent, cohort-specific, or not recovered by the profiling method?

Technical variation in faecal sample processing and downstream statistical modelling can alter measured community composition and the robustness of microbiome-disease associations [7, 10]. Among these analytical sources of uncertainty, taxonomic profiling is particularly important because microbiome biomarkers are not observed directly from sequencing reads but are inferred through computational workflows that determine which taxa are detected and how their abundances are estimated [11]. For disease-associated signals with small between-group effect sizes or low relative abundance, choices involving taxonomic assignment, reference-database composition, abundance estimation, and feature filtering may influence whether a candidate biomarker is recovered and retained for downstream analysis [9, 12]. Indeed, host-phenotype classification can retain similar performance when relative-abundance profiles are reduced to presence-absence information, suggesting that reliable taxon detection may itself be a major determinant of microbiome biomarker performance [13].

Most evaluations of taxonomic profilers use simulated communities, mock communities, or defined reference mixtures that provide controlled ground truth but do not fully capture the ecological and technical complexity of human stool metagenomes [14–19]. Conversely, clinical metagenomes preserve this complexity but usually lack ground truth, making it difficult to distinguish true biological absence from profiler non-detection, abundance distortion, or taxonomic misassignment.

Here, we developed an *in silico* spike-in framework to measure the analytical recovery of CRC-associated microbial signals in clinical stool metagenomic backgrounds. We embedded 10 consistently reported CRC-associated taxa [3–5], either individually or as a mixed community, into stool metagenomes from two CRC cohorts spanning control, adenoma, and CRC backgrounds and a series of low abundance fractions. We used two widely used and methodologically contrasting workflow-database configurations as experimental cases to test whether the same known signals remained detectable, quantitatively interpretable, and statistically recoverable across a complete biomarker-discovery pipeline. Kraken2/Bracken combines k-mer-based read classification with species-level abundance re-estimation [20, 21], whereas MetaPhlAn 4 estimates community composition using clade-specific marker genes [22]. These contrasting analytical strategies provided a practical setting in which to determine whether identical implanted signals could produce different recovery outcomes. The two workflow–database configurations were used as contrasting analytical case studies and were not evaluated to establish comparative superiority. Their purpose was to reveal how identical implanted signals can be recovered, distorted, or lost across an end-to-end biomarker-discovery pipeline.

Using this framework, we addressed four questions. First, under which abundance and clinical-background conditions are low-abundance CRC-associated spike-ins detected and quantitatively recovered by contrasting profiling configurations? Second, which CRC-associated taxa are consistently recoverable, and which remain analytically difficult across conditions? Third, which implanted taxa fail to emerge as significant features in downstream differential-abundance analyses, and which analytical characteristics are associated with this failure? Fourth, can spike-in-derived information reduce false-positive biomarker calls and identify abundance distortions sufficiently systematic to support future calibration? Together, these analyses were designed to determine when failure to recover a CRC-associated signal reflects analytical limitations rather than biological absence.

## 2 Results

### 2.1 Study framework and baseline profiler-dependent differences in CRC-associated taxa

We developed an *in silico* spike-in framework to evaluate the recovery of recurrently reported CRC-associated taxa in human stool metagenomes (Fig. 1). To preserve the ecological and technical complexity of clinical stool samples while introducing a quantitative ground truth, we implanted 10 CRC-associated taxa (Table 1) into preprocessed metagenomes, either individually or as a mixed ten-species community.

**Fig. 1:**
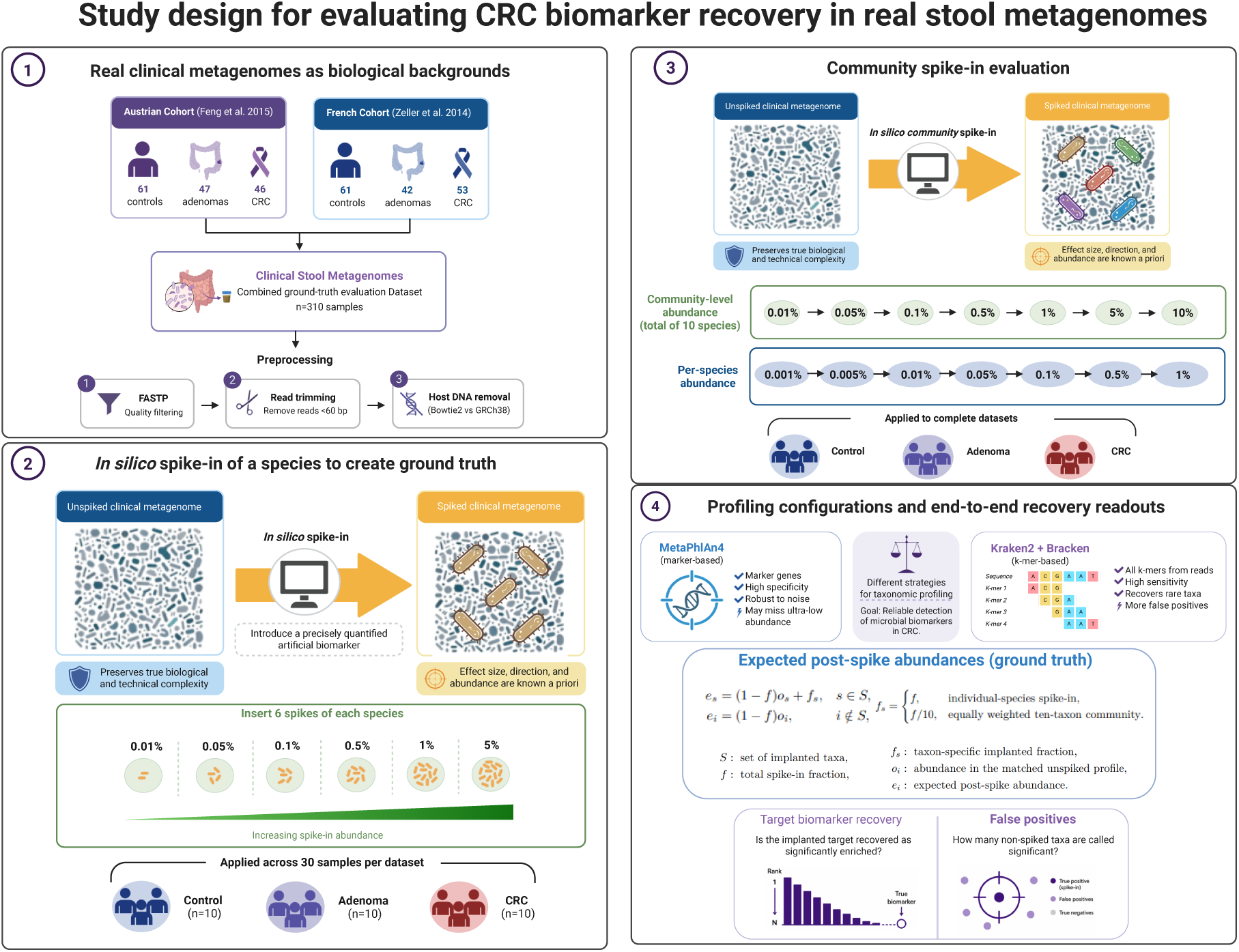
Study design for evaluating CRC biomarker recovery in human stool metagenomes. Human stool metagenomes from two public CRC cohorts were used as the biological baseline for *in silico* spike-in evaluation. CRC-associated taxa were introduced either individually or as a ten-species community at known abundance fractions. Original and spiked samples were profiled with MetaPhlAn 4 and Kraken2/Bracken, and analytical recovery was assessed using target detection, abundance accuracy, non-target abundance artefacts, and downstream differential-abundance recovery.

**Table 1:**
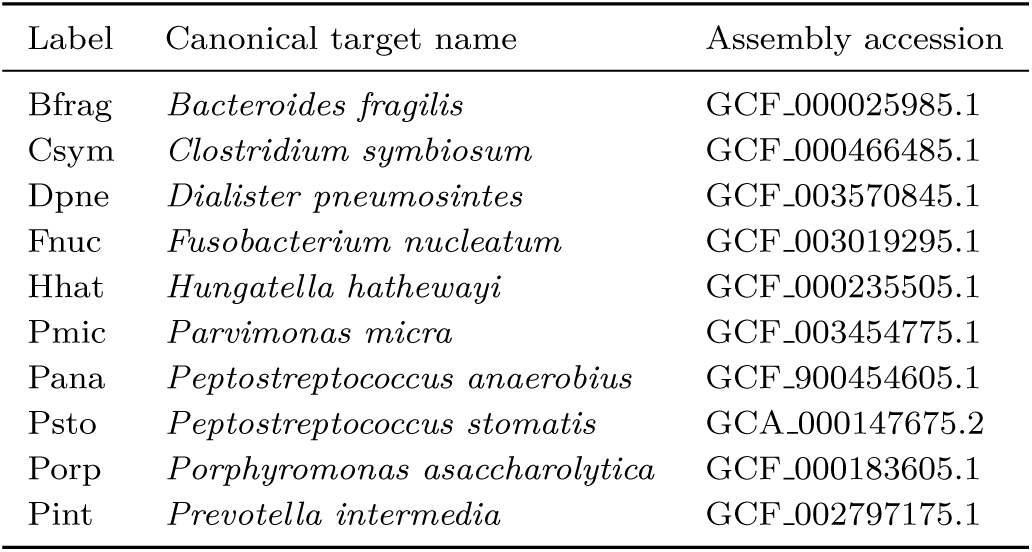
CRC-associated target taxa and reference genome assemblies used for the ground-truth spike-in evaluation. Taxa are reported using canonical species-level names. Profiler-specific taxonomic labels and aliases are provided in Supplementary Table A1.

The study comprised two complementary spike-in designs: an individual-species design, in which each target taxon was introduced separately, and a community design, in which all 10 taxa were introduced simultaneously. In the individual-species experiment, we selected 10 control, 10 adenoma, and 10 CRC samples from each of the FengQ 2015 and ZellerG 2014 cohorts, corresponding to 30 samples per cohort and 60 samples in total [1, 2]. Each target taxon was spiked independently into every selected sample at six final read-pair fractions: 0.01%, 0.05%, 0.1%, 0.5%, 1%, and 5%. These fractions covered a broad abundance range to allow us to evaluate recovery from very low to progressively stronger implanted signals. In the community experiment, we used all 154 samples from the FengQ 2015 cohort and all 156 samples from the ZellerG 2014 cohort, for a total of 310 samples, and introduced the same 10 taxa simultaneously as an equally weighted mixed community at seven total-community read-pair fractions: 0.01%, 0.05%, 0.1%, 0.5%, 1%, 5%, and 10%. Because each taxon contributed one tenth of the total community spike, the corresponding effective per-taxon fractions were 0.001%, 0.005%, 0.01%, 0.05%, 0.1%, 0.5%, and 1%.

Post-QC sequencing depth differed substantially between the two cohorts (Supplementary Fig. B1A). FengQ 2015 samples generally retained greater read-pair depth than ZellerG 2014 samples, whereas depth distributions overlapped broadly across diagnostic backgrounds within each cohort. Read-pair retention after preprocessing was high and relatively consistent in FengQ 2015, but was lower and more variable in ZellerG 2014, including a small number of samples with pronounced read loss (Supplementary Fig. B1B). Nevertheless, raw and post-QC sequencing depths were strongly concordant overall, indicating that preprocessing largely preserved the relative sequencing-depth structure of the original datasets (Supplementary Fig. B1C). Because spike-in levels were defined as fractions of the final read-pair count, variation in post-QC depth resulted in different absolute numbers of implanted read pairs at the same nominal spike fraction. We therefore examined whether post-QC sequencing depth was associated with target detection and quantitative recovery, particularly at the lowest spike-in levels.

Using this design, we evaluated recovery at three levels. First, we assessed whether implanted taxa were detected and quantified close to their expected abundances. Second, because adding spike-in reads increases the total number of reads, non-target taxa were expected to show only a proportional decrease in relative abundance. Deviations from this dilution-only expectation were considered abundance artefacts. Third, we tested whether the implanted taxa were recovered as significant features in downstream differential-abundance analyses. All original and spiked metagenomes were profiled with MetaPhlAn 4 and Kraken2/Bracken, allowing the same implanted signals to be followed through two contrasting analytical configurations under identical biological backgrounds.

All ten implanted genomes had a closely related UHGG v2.0.2 species representative, with Mash distances ranging from 0 to 0.0345, and every target species was associated by taxonomic label with at least one MetaPhlAn 4 vJan25 SGB (Supplementary Table A2). This confirmed species-level representation in both reference systems, although it does not imply equivalent compatibility between each spike-in genome and the workflow-specific marker or k-mer reference space.

Again, a key feature of this study is that spike-ins were added to the existing microbial communities of real human stool metagenomes, rather than to artificial communities lacking a biological background. We therefore first quantified the baseline prevalence and abundance of the implanted taxa in the unspiked metagenomes. Several implanted taxa were already present before spike-in, but their baseline prevalence differed markedly between profilers, cohorts, and diagnostic groups (Fig. 2A; Supplementary Fig. B2A). Kraken2/Bracken generally reported higher baseline prevalence than MetaPhlAn 4, particularly for *Fusobacterium nucleatum*, *Parvimonas micra* and *Prevotella intermedia*. In adenoma samples, these taxa were more frequently detected by Kraken2/Bracken, whereas MetaPhlAn 4 reported fewer positive baseline calls. These differences indicate that several candidate CRC biomarkers lie close to the effective detection limits of these taxonomic profilers, such that workflow choice can determine whether a baseline signal is available for downstream biomarker analysis.

**Fig. 2:**
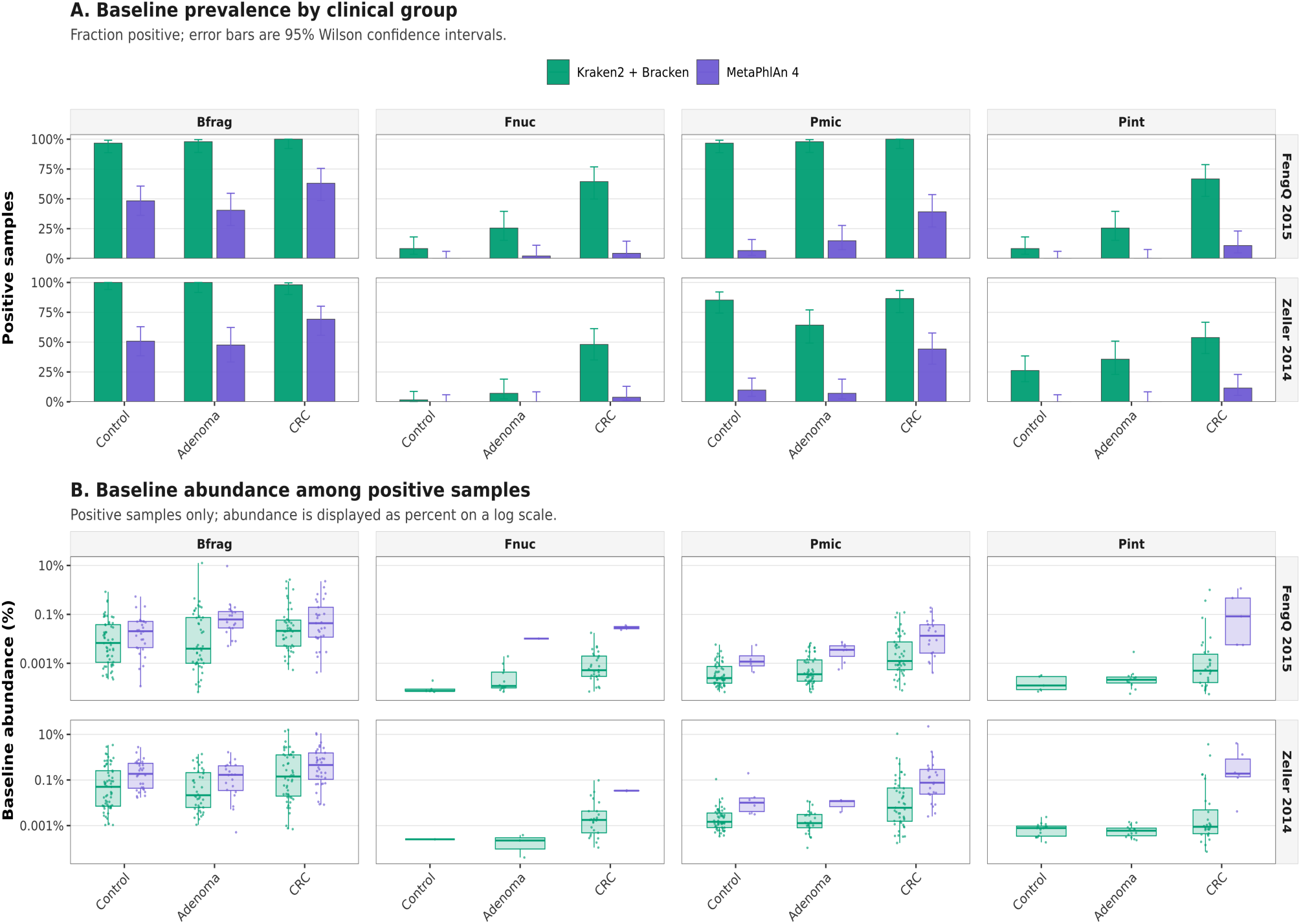
Profiler-dependent differences in baseline detection and abundance of CRC-associated taxa across full cohorts. **(A)** Full-cohort baseline prevalence of selected CRC-associated taxa across diagnostic groups in the FengQ 2015 and ZellerG 2014 cohorts. Bars show the percentage of samples with non-zero baseline abundance for each taxon, stratified by diagnostic group and profiler. Error bars indicate 95% Wilson confidence intervals. **(B)** Baseline abundance distributions among positive samples, defined as samples with abundance greater than zero for the corresponding profiler–taxon combination. Boxplots summarize the abundance distributions, and points represent individual samples. The abundance axis remains logarithmic, with tick labels expressed as percentages. Selected taxa are shown for clarity; the complete ten-taxon panel is provided in Supplementary Fig. B2.

Baseline abundance estimates also differed between profilers (Fig. 2B; Supplementary Fig. B2B). Among samples in which a target was detected, MetaPhlAn 4 generally assigned higher abundances than Kraken2/Bracken, particularly for *P. intermedia* and *P. micra*. Thus, the profilers differed both in whether taxa were detected and in the abundance assigned to positive detections. Kraken2/Bracken tended to report more frequent low-abundance detections, whereas MetaPhlAn 4 reported fewer but generally higher-abundance positive calls. Because expected post-spike abundances were calculated relative to these baseline profiles, this profiler-specific discordance directly influences the recovery of low-abundance implanted signals.

### 2.2 Individual-species spike-ins reveal workflow- and taxon-specific recovery limits at low abundance

We first evaluated the quantitative recovery of individually implanted CRC-associated taxa at the three lowest spike fractions (0.01%, 0.05%, and 0.10%; Fig. 3). Good quantitative recovery was defined more strictly than detection: a sample was classified as good only when the reported post-spike abundance was between 90% and 110% of the expected value, corresponding to an absolute relative recovery error ≤ 10%. Positive abundance estimates outside this interval were assigned to the intermediate or poor/missed classes according to their recovery error, while non-detections were classified as poor/missed. Figure 3A shows the percentage of samples meeting the good-recovery criterion.

**Fig. 3:**
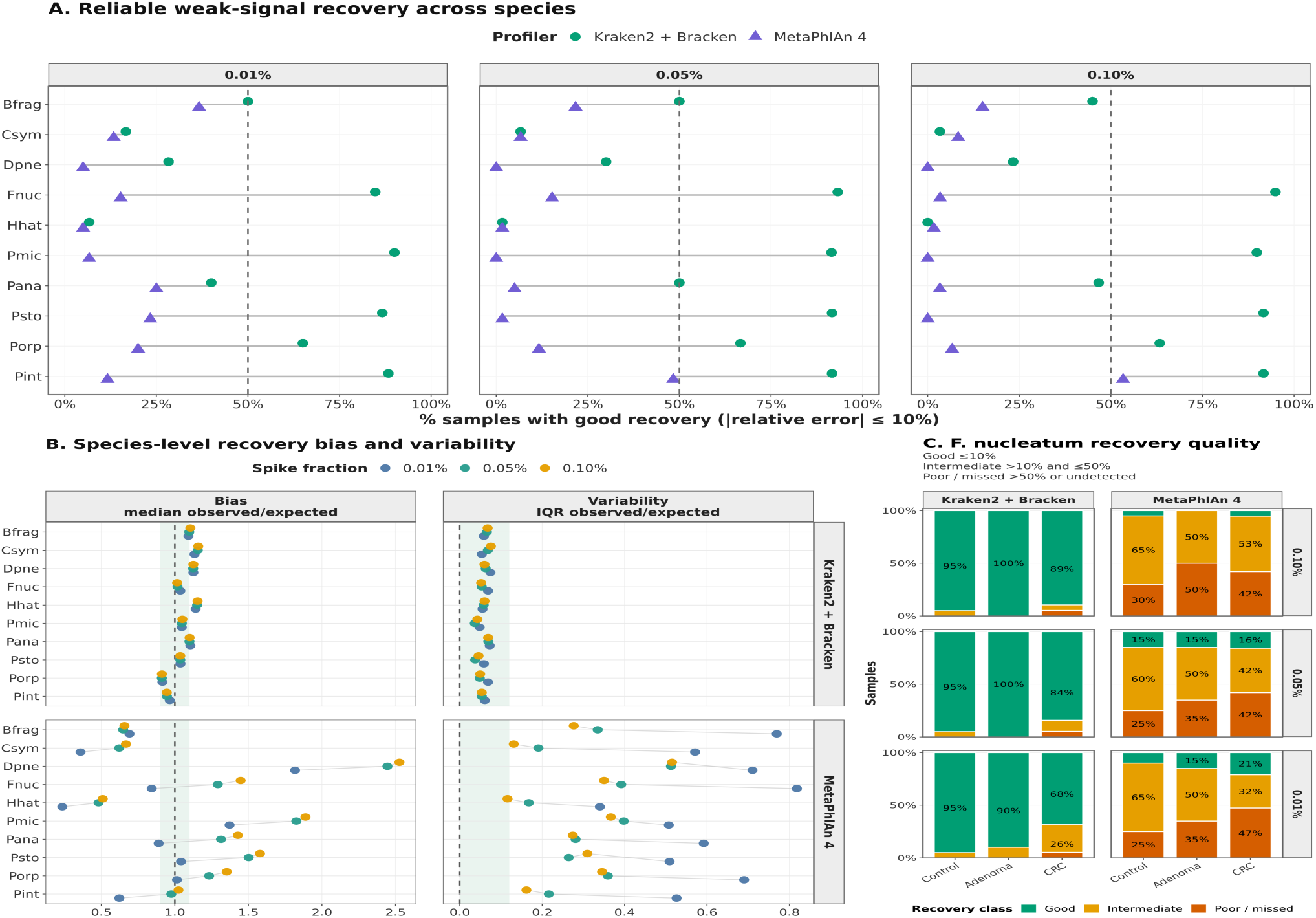
Individual-species spike-ins reveal workflow- and taxon-specific quantitative-recovery limits at low abundance. **(A)** Percentage of samples with Good quantitative recovery for each target taxon at the three lowest individual-species spike fractions (0.01%, 0.05%, and 0.10%). Good recovery required an absolute relative recovery error of at most 10%, equivalent to a reported abundance between 90% and 110% of the expected post-spike abundance. Points show the percentage of samples meeting this criterion for Kraken2/Bracken and MetaPhlAn 4. Grey lines connect the two profilers for the same taxon and spike fraction. The dashed vertical line marks 50% of samples. **(B)** Central bias and between-sample variability of quantitative recovery at the same spike fractions. Bias is summarized as the median observed-to-expected abundance ratio, for which a value of 1 indicates that estimates are centred on the expected abundance. Variability is summarized as the interquartile range of the observed-to-expected ratios. Points represent spike fractions, and grey lines connect fractions for the same taxon within each profiler. A median ratio close to 1 does not necessarily imply that most individual samples satisfy the stricter *±*10% Good-recovery criterion in panel A. **(C)** Recovery-class composition for individually spiked *Fusobacterium nucleatum* across diagnostic backgrounds and the three lowest spike fractions. *F. nucleatum* was selected as a clinically relevant example showing pronounced profiler-dependent recovery. Samples were classified as Good when the absolute relative recovery error was *≤* 10%, Intermediate when the error was *>* 10% and *≤* 50%, and Poor/missed when the error was *>* 50% or the target was not detected. All-fraction summaries are provided in Supplementary Figs. B3–B5.

At the 0.01% spike fraction, Kraken2/Bracken showed particularly high good-recovery rates for four taxa: *F. nucleatum*, *P. micra*, *P. stomatis*, and *P. intermedia*. For each of these taxa, approximately 85–90% of samples were accurately recovered (Fig. 3A; Supplementary Fig. B3). MetaPhlAn 4 showed substantially lower good-recovery rates for the same taxa. Good recovery remained very low for *F. nucleatum*, *P. micra*, and *P. stomatis*, while *P. intermedia* showed some improvement at higher spike fractions.

In contrast, some taxa were difficult to recover at the correct abundance regardless of the profiling workflow. *C. symbiosum* and *H. hathewayi* met the accurate-recovery criterion in fewer than 20% of samples under both Kraken2/Bracken and MetaPhlAn 4 across the low-abundance range (Fig. 3A). This indicates that the recoverability of an implanted signal depends not only on its abundance or the selected workflow, but also on the identity of the target taxon.

Panel B provides additional information about why a taxon may have a low good-recovery rate in panel A. The median observed-to-expected ratio shows whether the abundance estimates are centred on the expected value, while the IQR shows how much they vary among samples (Supplementary Fig. B5). Kraken2/Bracken estimates were generally centred close to the expected abundance and varied relatively little. *B. fragilis* provides a useful example. Its median ratio was close to one, indicating little overall bias, but only approximately half of the samples fell within the strict ±10% interval at the lowest spike fraction. In other words, the estimates were generally close to the expected value, but many were not close enough to be classified as accurately recovered. The lower percentage in panel A therefore reflects modest sample-level errors rather than substantial systematic overestimation or underestimation.

MetaPhlAn 4 showed a different pattern. For several taxa, the abundance estimates were shifted away from the expected value, more variable among samples, or both. *D. pneumosintes* and *P. micra* were generally overestimated, whereas *B. fragilis*, *C. symbiosum*, and *H. hathewayi* were generally underestimated. Several taxa also showed wide IQRs, indicating that the magnitude of the error differed considerably among samples. Therefore, failure to meet the accurate-recovery criterion can have several meanings: estimates may be consistently too high, consistently too low, highly variable, or absent altogether.

Recovery-class patterns also varied across clinical backgrounds, as illustrated using *F. nucleatum* in Fig. 3C and across all implanted taxa and fractions in Supplementary Fig. B4. *F. nucleatum* was selected because it is a recurrently reported CRC-associated taxon and showed one of the clearest workflow contrasts in panel A. Kraken2/Bracken produced high good-recovery rates across most backgrounds and spike fractions, ranging from 68.4% in the CRC background at 0.01% to 100% in the adenoma background at 0.05% and 0.10%. MetaPhlAn 4 showed substantially lower and more background-dependent good recovery. At 0.05% and 0.10%, many MetaPhlAn 4 profiles remained in the intermediate or poor/missed classes, showing that increasing the implanted abundance might have improved detectability but did not consistently result in good quantitative recovery.

These spike-in results provide a possible explanation for the lower baseline prevalence of *F. nucleatum* reported by MetaPhlAn 4. Some low-abundance baseline signals may have been missed because they fell within a range in which the workflow showed limited recovery. Conversely, the higher prevalence reported by Kraken2/Bracken may partly reflect greater sensitivity to low-abundance *F. nucleatum*, although the absence of ground truth in the unspiked samples prevents determining which baseline calls were correct. Thus, the same low-abundance *F. nucleatum* signal was more consistently quantified by Kraken2/Bracken, whereas MetaPhlAn 4 produced substantially more distorted and variable estimates.

Together, these results indicate distinct operating characteristics rather than a universal ranking of the two workflows. Kraken2/Bracken achieved more frequent good recovery for several taxa at low abundance, while MetaPhlAn 4 showed stronger taxon-specific bias and variability. Both workflows nevertheless struggled with targets such as *H. hathewayi* and *C. symbiosum*. Failure to recover a low-abundance microbial signal should therefore not be interpreted as biological absence without considering both detection and quantitative recovery by the selected workflow.

### 2.3 Recovery limits propagate to downstream biomarker detection

We next asked whether the quantitative-recovery limitations identified in Section 2.2 affected downstream biomarker detection. Here, biomarker recovery refers to the emergence of an implanted taxon as a significant group-level differential-abundance feature. It does not require successful quantitative recovery in every individual sample. Rather, the implanted signal must be recovered with sufficient consistency across the group to produce a detectable abundance shift. For each implanted taxon, profiler, cohort, and clinical background, we compared spiked and unspiked profiles and recorded the lowest spike fraction at which the implanted target was detected as significantly enriched. We refer to this value as the minimum biomarker-detection fraction. Lower values indicate that a weaker implanted signal was sufficient to produce a significant differential-abundance result, whereas NR indicates that the target was not significantly recovered at any tested spike fraction.

Clinical background strongly influenced the minimum biomarker-detection fraction (Fig. 4A). In control backgrounds, most targets were significantly recovered at the lowest tested fraction of 0.01%, particularly with Kraken2/Bracken. Recovery thresholds were also generally low in adenoma backgrounds, although several profiler-taxon combinations required stronger signals. In CRC backgrounds, the minimum required fractions were typically higher and more heterogeneous. This pattern is consistent with a fixed implanted increment being more difficult to distinguish when the corresponding taxon is already present at a higher or more variable baseline abundance.

**Fig. 4:**
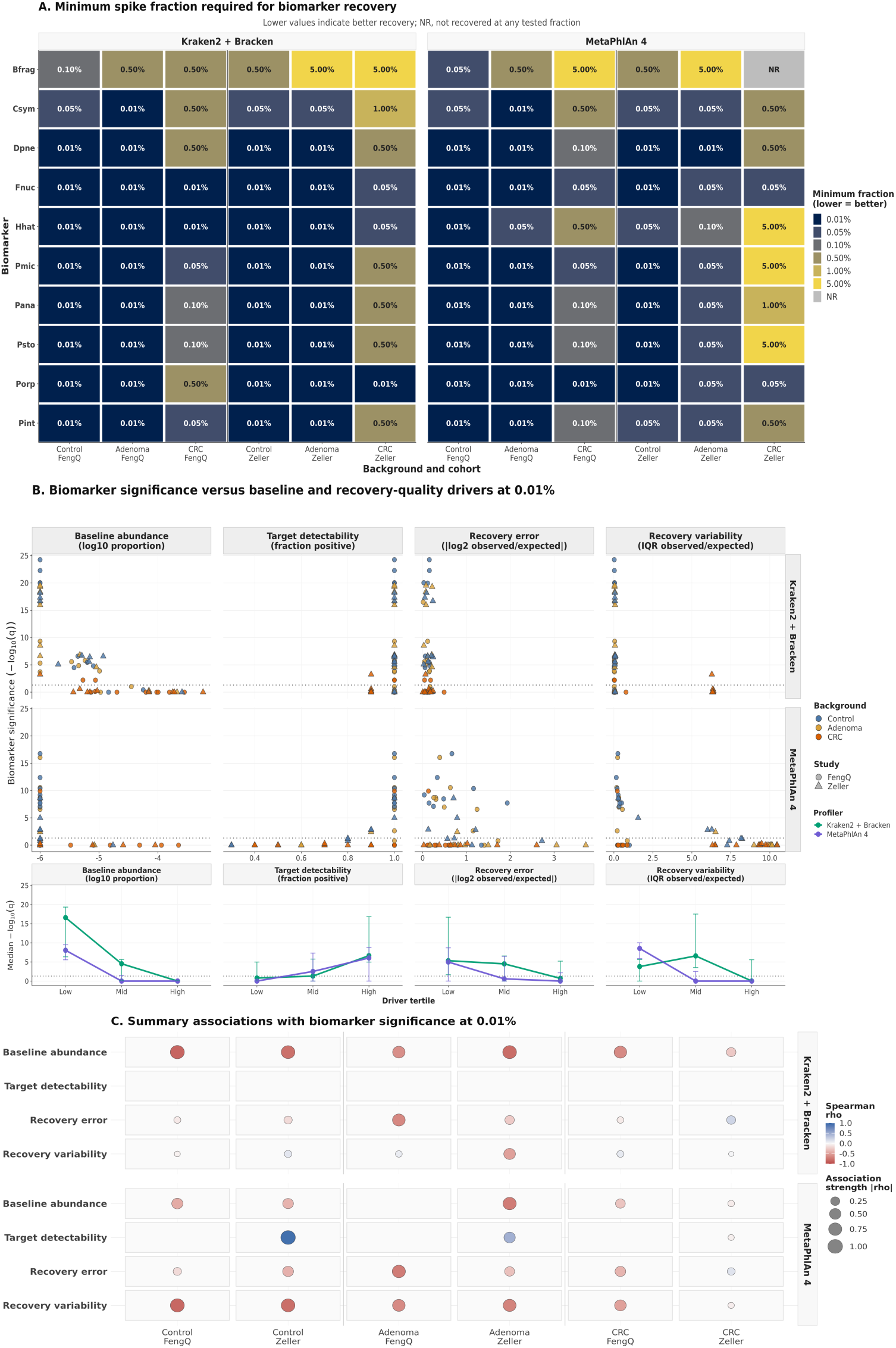
Biomarker recoverability depends on profiler, taxon, and biological background. **(A)** Minimum spike fraction required for each independently spiked taxon to be recovered as a significantly enriched biomarker in the spike-status differential-abundance recovery analysis. Rows represent target taxa, columns represent cohort and diagnostic-background combinations, and profilers are shown in separate facets. Tile colour and text report the minimum recovered spike fraction; lower values indicate recovery at a weaker implanted abundance. NR indicates that the target was not recovered at any tested spike fraction. **(B)** Relationship between biomarker significance at the weakest individual-species spike fraction, 0.01%, and four interpretable taxonomy-side drivers: baseline abundance, target detectability, recovery error, and recovery variability. Biomarker significance is expressed as *−* log_10_(*q*) for the expected target taxon. Points represent individual target–background combinations; point colour identifies diagnostic background and point shape identifies cohort. The lower panels show the median significance within driver tertiles, with error bars representing the interquartile range. **(C)** Spearman correlations between biomarker significance at 0.01% and the same four drivers, stratified by profiler, cohort, and diagnostic background. Circle colour indicates the direction and magnitude of Spearman’s *ρ*, and circle size indicates *|ρ|*. Positive correlations indicate that higher driver values are associated with stronger target-biomarker significance, whereas negative correlations indicate associations with weaker significance. Blank cells indicate combinations for which a correlation could not be estimated because of insufficient variation or too few valid observations.

The minimum spike fraction required for significant biomarker detection also differed substantially among target taxa (Fig. 4A). *B. fragilis* was the most consistently difficult target to recover as a biomarker. Depending on profiler, cohort, and clinical background, it required spike fractions between 0.05% and 5%, and it was not recovered at any tested fraction by MetaPhlAn 4 in the ZellerG 2014 CRC background. By contrast, *F. nucleatum* and *P. asaccharolytica* were recovered at relatively weak fractions across all evaluated contexts, generally at 0.01–0.05%. Several other targets showed pronounced background dependence. *D. pneumosintes* was recovered at 0.01% in control and adenoma backgrounds but generally required 0.10–0.50% in CRC backgrounds. *P. micra* and *P. stomatis* were recovered at 0.01–0.05% in most control and adenoma contexts, but required stronger signals in CRC backgrounds, reaching 5% for MetaPhlAn 4 in the ZellerG 2014 CRC cohort. A similarly strong context dependence was observed for *H. hathewayi*, which required 5% under MetaPhlAn 4 in the ZellerG 2014 CRC background. *P. intermedia* was generally recovered at low fractions but required up to 0.50% in the ZellerG 2014 CRC background. Thus, minimum biomarker-detection thresholds depended jointly on target identity, profiler, cohort, and clinical background.

We next examined whether baseline abundance and the quantitative-recovery properties measured in Section 2.2 were associated with biomarker significance at the weakest individual-species spike fraction, 0.01% (Fig. 4B). Identical implanted fractions produced a wide range of significance values, from effective non-detection to strong statistical evidence. Baseline abundance showed the clearest overall relationship: targets with higher baseline abundance generally exhibited weaker biomarker significance, consistent with a fixed implanted increment being more difficult to distinguish from a larger baseline signal.

Target detectability, recovery error, and recovery variability were also associated with biomarker significance in some contexts. More frequent target detection generally resulted in stronger significance, whereas larger recovery errors and greater between-sample variability tended to result in weaker significance. However, these relationships were not uniform across profilers, cohorts, and clinical backgrounds (Fig. 4C). Blank cells indicate combinations for which a correlation could not be estimated because the corresponding driver showed insufficient variation.

Together, these results show that downstream biomarker detection depends partly on whether an implanted abundance increment is detected and quantified accurately and consistently. However, quantitative recovery alone does not determine statistical significance. Baseline target abundance, clinical background, cohort, profiler, and target identity jointly determine whether the same implanted signal emerges as a significant biomarker.

### 2.4 Community spike-ins reproduce recovery patterns and reveal a sensitivity–specificity tradeoff

We next asked whether the recovery patterns observed in the individual-species experiment were preserved when all 10 CRC-associated taxa were introduced simultaneously as a mixed community. Because each taxon contributed one tenth of the total community spike, total community spike fractions were converted to effective per-taxon fractions to enable direct comparison between the two spike-in designs.

Community spike-ins broadly reproduced the taxon-specific good-recovery patterns observed in the individual-species experiment (Fig. 5A). With Kraken2/Bracken, highly recoverable taxa such as *F. nucleatum*, *P. intermedia*, *P. micra*, and *P. stomatis* remained among the best recovered at matched effective fractions. In contrast, *D. pneumosintes*, *C. symbiosum*, and *H. hathewayi* remained among the most difficult taxa to recover. MetaPhlAn 4 again showed lower good-recovery rates for several taxa, particularly at the lowest matched fractions. Thus, the workflow- and taxon-specific patterns identified using individual-species spike-ins were largely preserved when the targets were implanted together.

**Fig. 5:**
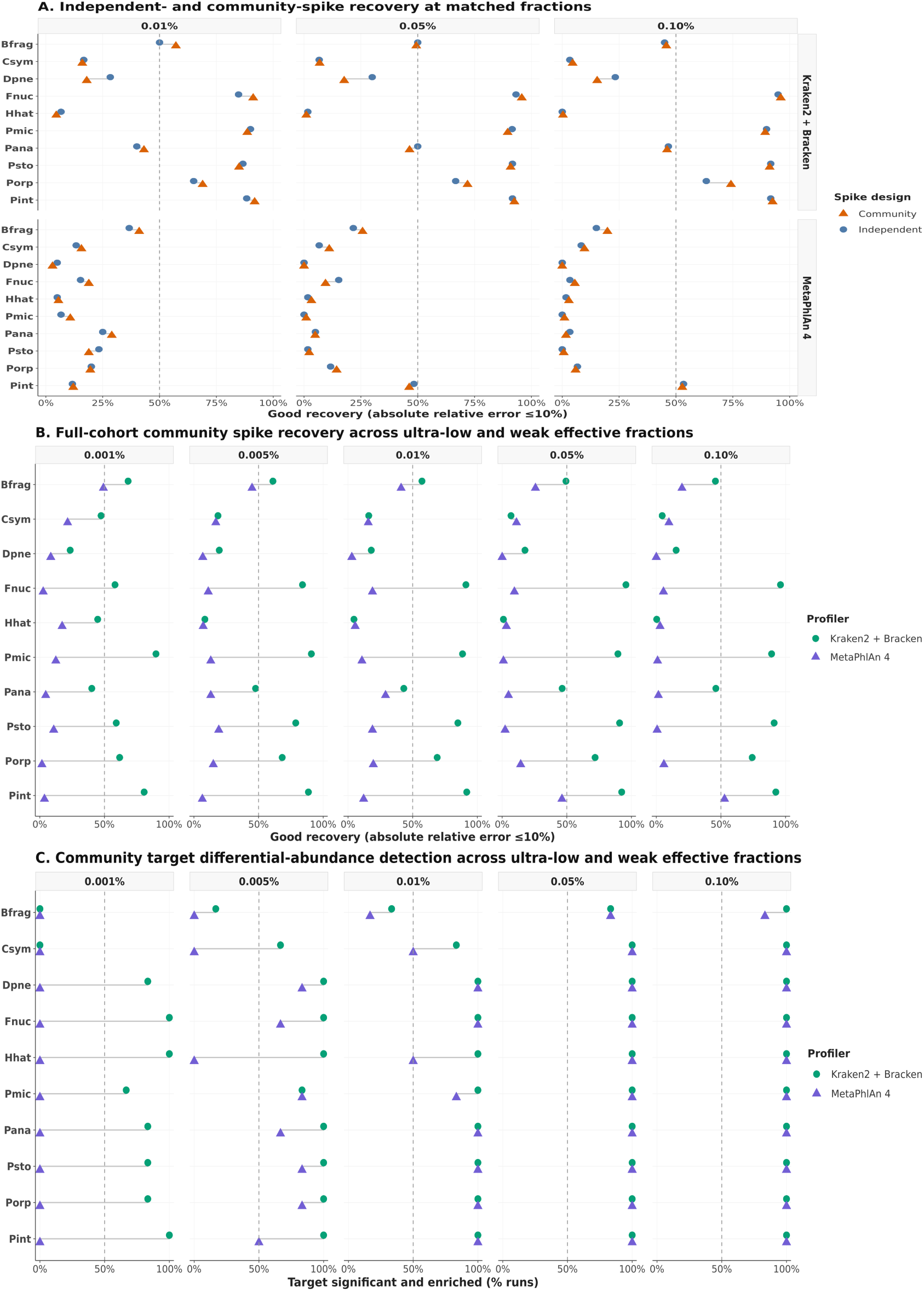
Community spike-ins preserve individual-species recovery patterns and downstream biomarker detection. **(A)** Comparison of individual-species and community spike-in recovery at matched effective per-taxon fractions. Points show the percentage of samples with Good quantitative recovery for each target taxon, profiler, and spike-in design. Good recovery required an absolute relative recovery error of at most 10%. Community spike fractions were converted to effective per-taxon fractions by dividing the total community fraction by the ten implanted taxa. The dashed vertical line marks Good recovery in 50% of samples. **(B)** Full-cohort community spike-in recovery across ultra-low and weak effective per-taxon fractions. Points show the percentage of samples with Good recovery for each implanted taxon and profiler. The dashed vertical line marks 50% of samples. **(C)** Full-cohort community-target detection in downstream differential-abundance analyses. Points show the percentage of runs in which each implanted community member was detected as significantly enriched in spiked samples. Results are stratified by effective per-taxon fraction and profiler, and the dashed vertical line marks detection in 50% of runs. Corresponding all-fraction recovery and differential-abundance summaries are provided in Supplementary Figs. B7 and B11.

The full-cohort community experiment extended this analysis to effective per-taxon fractions as low as 0.001% (Fig. 5B). At this fraction, Kraken2/Bracken achieved good recovery for many taxa in more than half of the analysed samples, whereas most MetaPhlAn 4 targets again showed good recovery in only a small proportion of samples. Taxa that were comparatively easy or difficult to recover in the individual-species experiment generally showed the same pattern in the community experiment.

Across the complete effective-fraction series, Kraken2/Bracken observations were predominantly assigned to the good or intermediate classes, with relatively few poor/missed recoveries. MetaPhlAn 4 showed a high proportion of poor/missed recoveries at the lowest effective fractions. As the implanted fraction increased, many MetaPhlAn 4 observations shifted from poor/missed to intermediate rather than consistently reaching good recovery (Supplementary Fig. B6). These transitions differed among implanted taxa (Supplementary Figs. B7–B9). Greater implanted read support was likewise associated with fewer poor/missed MetaPhlAn 4 recoveries, but mainly increased the proportion classified as intermediate rather than good (Supplementary Fig. B10). Kraken2/Bracken recovery-class composition changed comparatively little across read-support tertiles.

We next tested whether the implanted community members were detected as significantly enriched targets in downstream differential-abundance analyses (Fig. 5C). This analysis addressed a different outcome from panels A and B: panels A and B measure quantitative abundance accuracy, whereas panel C measures whether the implanted target emerged as statistically significant. Kraken2/Bracken detected many targets as significantly enriched even at the 0.001% effective fraction, and most targets approached complete detection by 0.005–0.01%. MetaPhlAn 4 showed little target detection at 0.001%, mixed detection at 0.005%, and substantially higher detection for many, but not all, targets by 0.01%. The complete fraction series showed the same abundance-dependent pattern, with Kraken2/Bracken detecting many implanted targets at lower effective fractions than MetaPhlAn 4 (Supplementary Fig. B11).

The greater target sensitivity of Kraken2/Bracken was accompanied by a substantial off-target differential-abundance burden (Supplementary Fig. B12). The number of significantly enriched non-target features increased with effective spike fraction for Kraken2/Bracken, whereas MetaPhlAn 4 maintained an off-target burden close to zero across the same range. Thus, Kraken2/Bracken detected weaker implanted targets more readily but generated a considerably noisier differential-abundance background. MetaPhlAn 4 provided greater specificity but showed weaker target detection and less accurate quantitative recovery for several low-abundance taxa.

Together, the community experiment reproduced the main workflow- and taxon-specific recovery patterns observed with individual-species spike-ins and showed that these patterns propagated to downstream biomarker detection. It also revealed that neither workflow was uniformly preferable: Kraken2/Bracken provided greater sensitivity to low-abundance targets at the cost of more off-target discoveries, whereas MetaPhlAn 4 provided a cleaner differential-abundance background at the cost of reduced recovery of several weak signals.

### 2.5 Spike-derived artefact exclusion reduces off-target differential-abundance calls and partially transfers between cohorts

The community spike-in experiment allowed us to test whether enriched off-target differential-abundance calls were associated with deviations from the dilution-only expectation. Because only the 10 implanted taxa were expected to increase after spike-in, all other taxa should change only through compositional dilution. For each workflow and spike fraction, a non-target taxon was classified as artefact-prone when either its mean absolute relative error or the standard deviation of its relative error exceeded 5%.

Enriched off-target differential-abundance calls were strongly concentrated among artefact-prone taxa (Fig. 6A). Under Kraken2/Bracken, 54.9% of non-target taxa were classified as artefact-prone, compared with only 0.5% under MetaPhlAn 4. Among taxa identified as enriched off-target differential-abundance features, these proportions increased to 96.3% and 100%, respectively. Thus, off-target discoveries were concentrated among taxa showing abnormal abundance behaviour rather than being distributed randomly across the workflow-derived feature tables. The same pattern was observed across the broader community spike-fraction series (Supplementary Fig. B13).

**Fig. 6:**
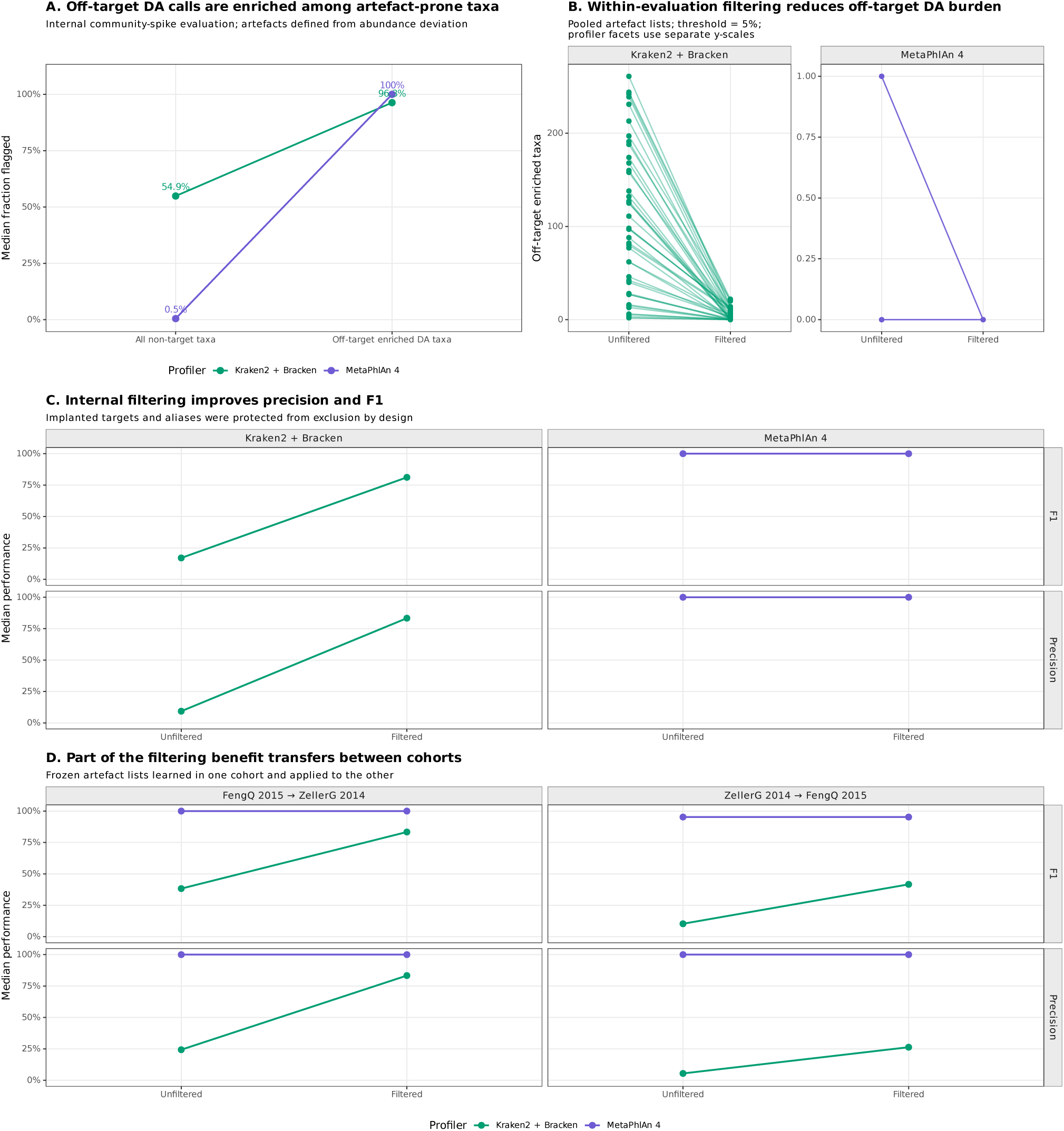
Spike-derived artefact exclusion reduces off-target differential-abundance calls and partially transfers between cohorts. **(A)** Enrichment of artefact-prone taxa among significantly enriched off-target differential-abundance calls in the internal community spike-in evaluation. For each profiler and analysis context, a non-target taxon was classified as artefact-prone when either its mean absolute relative error or the standard deviation of its relative error from the dilution-only expectation exceeded 5%. Points show the median fraction classified as artefact-prone among all non-target taxa and among enriched off-target differential-abundance taxa, pooled across cohort, diagnostic-background, and spike-fraction contexts. Percentage labels give the corresponding median fractions. **(B)** Within-evaluation exclusion using pooled profiler-specific artefact lists derived from the community spike-in data across both cohorts. Lines connect the unfiltered and filtered numbers of enriched off-target taxa within the same analysis context. Profiler facets use separate y-axis scales. **(C)** Median differential-abundance-list precision and F1 score before and after internal artefact-aware exclusion. The ten implanted targets and all mapped profiler-specific aliases were protected from exclusion by design. **(D)** Reciprocal cross-cohort transfer of artefact-aware exclusion. Frozen profiler-specific artefact lists were learned in one cohort and applied without modification to significant differential-abundance calls in the other cohort. Panels show median precision and F1 score before and after exclusion. Colours indicate profiler.

Beyond their overall burden, enriched off-target differential-abundance calls showed marked taxon-specific structure (Supplementary Fig. B14). For Kraken2/Bracken, the identities and recurrence of off-target features in the individual-species experiments depended on the implanted taxon, and same-genus taxa were overrepresented among enriched off-target calls for several targets (Supplementary Fig. B14A,B). MetaPhlAn 4 produced no enriched off-target calls in the individual-species analyses. In the community experiment, recurrent off-target calls were concentrated among taxa showing larger deviations from the dilution-only abundance expectation (Supplementary Fig. B14C). Some Kraken2/Bracken off-target features recurred across both individual-species and community-spike designs, although the largest recurrent set was specific to the community experiment. These findings indicate that the off-target burden was not randomly distributed across taxa, but reflected reproducible and partly target-dependent profiler behaviour.

We next tested whether this information could be used to reduce the off-target burden. Workflow-specific artefact lists were constructed from the pooled community spike-in data across both cohorts and applied to the corresponding differential-abundance results. The 10 implanted targets and their mapped aliases were protected from exclusion by design. This internal exclusion strongly reduced the number of enriched off-target taxa, particularly for Kraken2/Bracken, for which some analysis contexts initially contained tens to hundreds of off-target features (Fig. 6B). Because off-target taxa were removed while implanted targets were retained, differential-abundance-list precision and F1 score increased substantially (Fig. 6C). MetaPhlAn 4 already had high unfiltered precision and F1 because its off-target burden was close to zero, leaving comparatively little room for improvement.

We then tested whether artefact information transferred between cohorts. Artefact lists learned in one cohort were applied without modification to significant differential-abundance calls in the other cohort (Fig. 6D). For Kraken2/Bracken, cross-cohort exclusion improved precision and F1 in both directions, although the improvement was greater when the list learned in FengQ 2015 was applied to ZellerG 2014 than in the reciprocal direction. MetaPhlAn 4 retained high performance before and after exclusion because few off-target calls were present initially.

Together, these results show that spike-derived abundance deviations can identify taxa that disproportionately contribute to enriched off-target differential-abundance calls. The strongest improvements were obtained when artefact lists were applied within the same spike-in evaluation from which they were derived. Nevertheless, partial transfer between cohorts indicates that some workflow-induced abundance artefacts are reproducible.

### 2.6 Spike-in abundance-response modelling reveals workflow- and taxon-specific calibratability

We next asked whether the abundance distortions observed in the community spike-in experiment were sufficiently systematic to support quantitative calibration. Expected post-spike abundance was modelled separately for each workflow–taxon combination using linear models and generalized additive models with grouped cross-validation.

Expected and observed post-spike abundances showed strong but workflow- and taxon-specific relationships (Fig. 7A). For Kraken2/Bracken, the fitted abundance-response curves were generally close to the identity expectation for the representative taxa shown, consistent with the comparatively small quantitative errors observed in the preceding recovery analyses. MetaPhlAn 4 showed larger differences from the identity expectation, including systematic overestimation, underestimation, and greater dispersion at low expected abundances. Nevertheless, these differences were generally structured rather than random, indicating that some abundance distortions may be predictable.

**Fig. 7:**
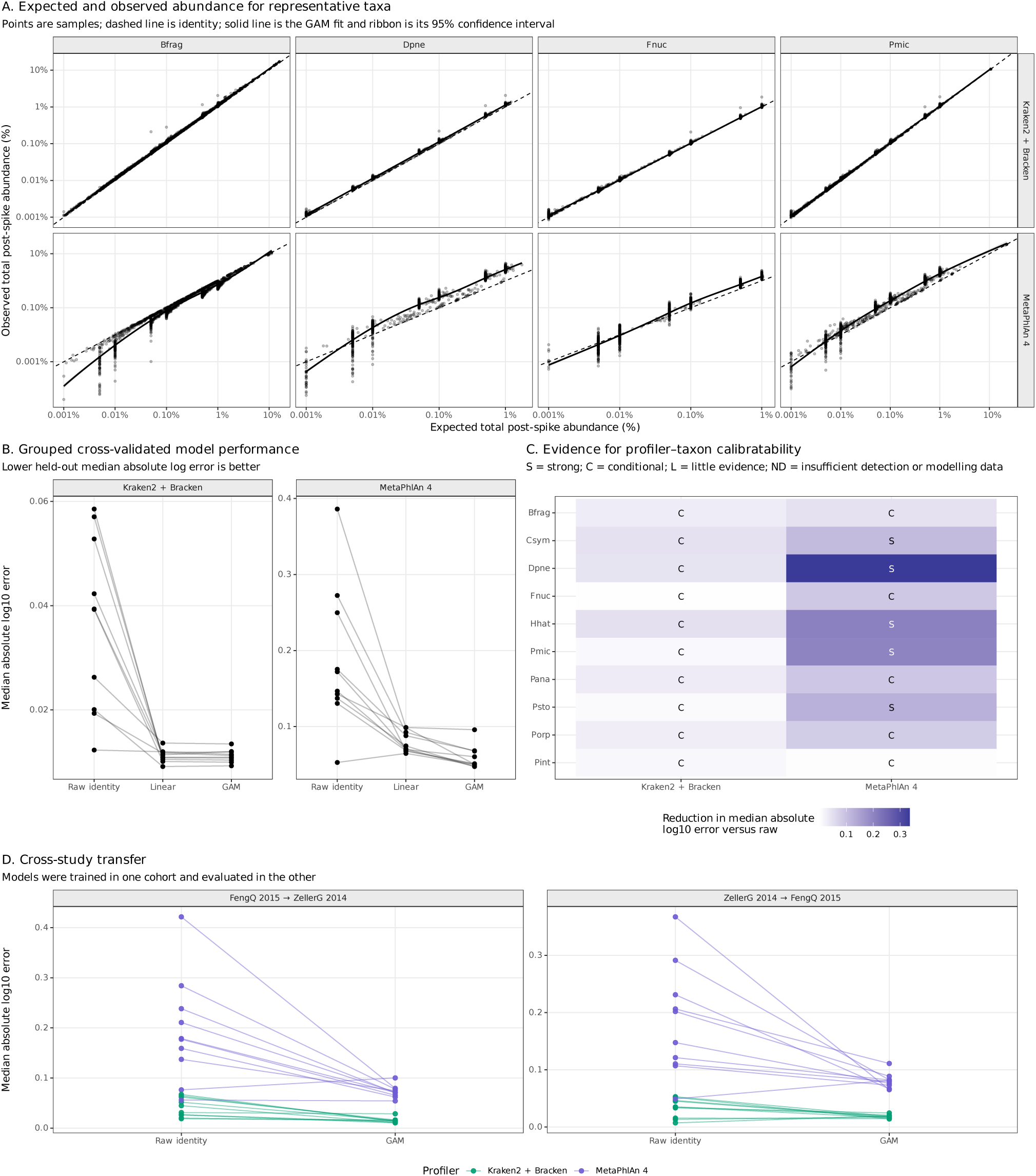
Spike-in abundance-response relationships reveal workflow- and taxon-specific calibratability. **(A)** Expected and observed post-spike abundances for representative community spike-in targets. Points represent individual samples, dashed lines indicate the identity expectation, and solid curves show generalized additive model (GAM) fits. Shaded ribbons indicate 95% confidence intervals for the fitted curves. Expected and observed abundances are displayed as percentages on logarithmic axes. **(B)** Grouped cross-validated prediction error for the raw identity expectation, linear regression, and inverse GAM. Lines connect model results for the same profiler–taxon combination. Lower held-out median absolute log_10_ error indicates better predictive performance. **(C)** Reduction in held-out median absolute log_10_ error obtained with the inverse GAM relative to the raw identity prediction. Tile colour represents the magnitude of the error reduction. Letters indicate Strong (S), Conditional (C), or Little (L) evidence of calibratability, or insufficient detection or modelling data (ND), according to the criteria defined in Methods. **(D)** Cross-cohort transfer of inverse GAMs. Models were trained in one cohort and evaluated in the other. Lines connect raw-identity and GAM-based errors for the same profiler–taxon combination, and colours indicate profiler. Lower values indicate better transferred prediction.

Grouped cross-validation showed that learning an abundance-response relationship generally improved prediction of expected abundance in held-out biological samples (Fig. 7B). Both linear models and inverse generalized additive models reduced median absolute log-scale error relative to using the raw workflow-reported abundance directly. For Kraken2/Bracken, the two model types performed similarly for most taxa, suggesting that relatively simple abundance-response relationships were often sufficient. For MetaPhlAn 4, generalized additive models provided additional improvement for several taxa, consistent with the stronger nonlinear departures observed in panel A.

The magnitude of improvement differed among workflow–taxon combinations (Fig. 7C). Several MetaPhlAn 4 combinations showed strong evidence of calibratability because the generalized additive model substantially reduced held-out prediction error. Other combinations showed only conditional evidence because the improvement was modest, the uncorrected error was already small, or recovery remained affected by non-detection and residual sample-to-sample variability. Kraken2/Bracken generally showed smaller absolute improvements because its uncorrected abundance estimates were already closer to the expected scale for many taxa.

The learned abundance-response relationships also showed partial transfer between cohorts (Fig. 7D). Models trained in FengQ 2015 generally reduced prediction error when evaluated in ZellerG 2014, and models trained in ZellerG 2014 showed a similar improvement when evaluated in FengQ 2015. However, residual error remained heterogeneous among workflow–taxon combinations, indicating that transferability was incomplete and that future calibration approaches will need to account for workflow, taxon, abundance range, and biological background.

Together, these results show that many workflow-specific abundance distortions contain a systematic component that can be modelled quantitatively. Calibration cannot recover taxa that were not detected and does not fully remove sample-specific error. Nevertheless, the observed abundance-response relationships provide a basis for future workflow- and taxon-specific abundance correction.

## 3 Discussion

Microbiome biomarkers are often interpreted as direct biological observations, yet they first pass through computational profiling workflows that determine whether candidate taxa are detected, how their abundances are estimated, and whether they ultimately become statistically significant features. By embedding known CRC-associated microbial signals into real stool metagenomes, we established a ground-truth framework for measuring these analytical effects under realistic biological backgrounds. Rather than asking only whether profilers classify taxa correctly, our approach tests whether weak microbial signals remain detectable, quantitatively interpretable, and statistically recoverable throughout a complete biomarker-discovery workflow. Across two independent CRC cohorts, analytical recoverability depended on the implanted taxon, profiling workflow, abundance regime, and clinical background. The same framework also showed that profiler-specific distortions can be used to identify off-target artefacts and to determine when quantitative correction may be feasible.

A central finding was that truly present weak microbial signals can fail to remain detectable, quantitatively accurate, or statistically recoverable across a complete biomarker-discovery pipeline. The two evaluated workflow-database configurations illustrated different analytical routes to this outcome. Kraken2/Bracken recovered several implanted CRC-associated taxa, including *F. nucleatum*, *P. micra*, *P. stomatis*, and *P. intermedia*, at lower abundance and generally with greater quantitative confidence than MetaPhlAn 4. In contrast, MetaPhlAn 4 produced substantially fewer enriched off-target features and therefore a cleaner background for downstream analysis. These results do not support a universal ranking of profilers. Instead, they show that improved recovery of weak targets can be accompanied by a greater burden of off-target discoveries, whereas a more conservative profile can reduce off-target calls while failing to recover or accurately quantify some weak signals. Thus, analytical non-recovery can arise through different workflow-dependent mechanisms, and negative biomarker findings should be interpreted in relation to the operating characteristics of the selected workflow.

This tradeoff is particularly relevant to early colorectal adenomas. Large metagenomic studies consistently distinguish CRC from healthy controls, whereas adenoma-associated microbial changes are weaker and less reproducible across cohorts [1–5]. Our findings complement these biological observations by showing that analytical recoverability becomes limiting in low-abundance regimes. The behaviour of *F. nucleatum* illustrates this point: recovery of identical implanted signals varied between profilers and across clinical backgrounds despite identical spike fractions. MetaPhlAn 4 showed lower and more heterogeneous quantitative recovery across the clinical backgrounds examined, whereas Kraken2/Bracken recovered the same abundance increments more consistently. These observations do not establish *F. nucleatum* as an adenoma biomarker, but they show that negative or inconsistent findings cannot automatically be interpreted as biological absence. Some discrepancies between studies may instead be caused because weak disease-associated signals fall close to the effective detection or quantification limits of the profiling workflow.

More broadly, our results establish a direct link between taxonomic recovery and downstream biomarker discovery. Previous taxonomic evaluations have primarily measured precision, recall, and abundance accuracy using simulated or mock communities [14, 15]. These metrics are essential, but they do not directly show whether weak microbial signals will enter a biomarker list after association testing. In our evaluation, target detectability, quantitative accuracy, and recovery stability all influenced biomarker significance. Taxa that were consistently detected and accurately quantified were more likely to be recovered as enriched biomarkers, whereas recovery error and sample-to-sample variability reduced statistical significance despite identical implanted abundance. This provides a mechanistic explanation for why the same microbial candidate may emerge as a biomarker in one study yet remain undetected in another.

The comparison between independent and community spike-ins shows that this ground-truth logic can be scaled beyond individual taxa. Independent spike-ins provide the clearest taxon-specific assessment because each organism is evaluated separately, but the design becomes increasingly impractical for larger biomarker panels. Community spike-ins largely preserved the recovery hierarchy observed in the independent experiments while allowing multiple candidate biomarkers to be evaluated simultaneously. The two designs are therefore complementary: independent spike-ins provide high-resolution validation of taxon-specific limits, whereas community spike-ins offer a scalable strategy for testing realistic multi-taxon signatures.

The ground-truth spike-in evaluation also provides information beyond target recovery. Because non-target taxa should change only through compositional dilution after spike insertion, deviations from this expectation identify profiler-induced abundance distortions. These artefact-prone taxa contributed disproportionately to enriched off-target differential-abundance calls, particularly for Kraken2/Bracken. Artefact-aware exclusion substantially improved differential-abundance-list precision while preserving implanted targets by design, and part of this benefit transferred between cohorts. In parallel, abundance-response modelling showed that many profiler-specific distortions were systematic rather than random. For several profiler-taxon combinations, expected and observed abundances followed reproducible relationships, and both linear and generalized additive models reduced prediction error relative to the original profiler estimates. These two applications address complementary problems: artefact-aware exclusion improves confidence in downstream biomarker lists, whereas abundance-response modelling tests whether detected signals can be interpreted on a more accurate quantitative scale.

These corrective analyses should nevertheless be viewed as proof of principle. Artefact-aware exclusion was strongest when applied within the same spike-in evaluation from which the artefact profiles were derived, and cross-cohort transfer was limited. Likewise, abundance-response relationships remained profiler-, taxon-, and cohort-specific and could not recover taxa that were not detected in the first place. The value of calibration is therefore not that it guarantees correction of every abundance estimate, but that it identifies conditions under which distortion is sufficiently systematic to be modelled. More extensive validation will be required before spike-derived artefact profiles or abundance-response functions can be used as general preprocessing resources.

Analytical recoverability was also strongly taxon-specific. Some recurrent CRC-associated taxa were recovered consistently at low abundance, whereas others remained difficult even when their implanted abundance was known. *C. symbiosum* and *H. hathewayi*, for example, were poorly recovered by both workflows, while *D. pneumosintes* showed both missed detection and substantial abundance distortion. Candidate biomarkers should therefore be evaluated not only for biological relevance but also for analytical recoverability within the abundance range at which they are expected to occur. These are distinct properties: a readily recoverable taxon is not necessarily biologically important, and a biologically important taxon is not necessarily analytically recoverable. Ground-truth spike-ins quantify analytical recoverability without making claims about biological causality.

This study has several limitations. We evaluated two public CRC cohorts and ten recurrently reported CRC-associated taxa, and additional testing across populations, sequencing depths, extraction protocols, taxa, and biomarker panels will be required to assess the generalizability of the observed recovery patterns. We selected two widely used but conceptually distinct workflow–database configurations as contrasting experimental cases. They were not intended to represent the full range of available taxonomic profilers. In addition, *in silico* spike-ins preserve realistic microbial community structure while providing exact abundance ground truth, but they do not reproduce all biological and technical sources of variation encountered in clinical studies, including DNA extraction bias, microbial growth dynamics, ecological interactions, host effects, and naturally occurring strain diversity. This limitation is particularly important because each implanted taxon was represented by a single reference assembly. Although all ten target species were represented in the evaluated reference systems (Supplementary Table A2), naturally occurring strains may differ from the implanted assemblies in marker-gene content, k-mer composition, and genomic similarity to the database representatives. Finally, the artefact-filtering and calibratability analyses remain specific to the workflows, databases, taxa, cohorts, and spike-in design evaluated here and require independent validation before standard use.

Despite these limitations, the framework addresses a fundamental problem in microbiome biomarker research: clinical metagenomes rarely provide ground truth. Embedding known microbial signals into real stool metagenomes helps distinguish true biological absence from cohort-specific variation and analytical non-recovery. This distinction is especially important for small disease-associated signals, including those reported for adenoma, because possible abundance changes may occur close to current analytical detection limits. Before concluding that a candidate biomarker is absent, irreproducible, or cohort-specific, it is therefore necessary to establish whether the analytical workflow can recover that signal confidently under realistic metagenomic conditions.

In conclusion, microbiome biomarker discovery is constrained not only by biological variability but also by profiler-specific analytical recoverability. The ground-truth spike-in evaluation revealed profiler-, taxon-, abundance-, and background-specific operating regimes that directly influenced downstream differential-abundance analyses. It also showed that profiler-induced distortions can be used to improve downstream interpretation through artefact-aware exclusion and future quantitative calibration. More broadly, this work establishes spike-in evaluation as a practical framework for determining whether candidate microbiome biomarkers are detectable, quantitatively interpretable, and statistically recoverable before biological conclusions are drawn. Incorporating ground-truth spike-in evaluations into microbiome biomarker studies should improve the interpretation of negative findings, motivate explicit validation of signal recoverability, and support more reliable translation of shotgun metagenomics into clinically useful biomarkers.

## 4 Methods

### 4.1 Study design

A ground-truth spike-in evaluation framework was developed to determine whether low-abundance colorectal cancer (CRC)-associated microbial signatures can be reliably detected when introduced into human stool metagenomic samples. Rather than evaluating taxonomic recovery only in mock communities or fully simulated samples, as in many taxonomic-profiling evaluations [14, 15], 10 known *in silico* microbial signals were added to preprocessed metagenomes from two public CRC cohorts. This design preserved endogenous community structure, host-associated background complexity, and cohort-level variation while providing known target identities and expected post-spike abundances.

Two publicly available CRC metagenomic cohorts were used: the Austrian cohort from Feng et al., comprising 61 controls, 47 adenomas, and 46 CRC cases [2], and the French cohort from Zeller et al., comprising 61 controls, 42 adenomas, and 53 CRC cases [1]. These cohorts were selected because they are widely used in CRC microbiome research, include control, adenoma, and CRC samples, and represent a weak-signal setting in which adenoma-associated microbial shifts are expected to be small relative to between-sample variability [1, 2, 5].

The two cohorts were used as independent biological backgrounds rather than as a pooled clinical discovery dataset. This design allowed recovery patterns to be evaluated across distinct studies while preserving study-specific microbial community structure, sequencing characteristics, and diagnostic-group composition. Generalizability was assessed by evaluating the consistency and transferability of recovery patterns, artefact signatures, and abundance-response relationships between cohorts.

The study comprised two complementary spike-in designs. In the individual-species experiment, a balanced subset of 10 samples per clinical group was selected from each cohort, yielding 30 samples per cohort and 60 samples in total. Samples were selected without formal randomization, matching, or stratification. The selected sample accessions are provided in Supplementary Table A3. Each selected sample was spiked independently with each of the 10 target taxa at six final read-pair fractions: 0.01%, 0.05%, 0.1%, 0.5%, 1%, and 5%. This design provided taxon-specific ground truth for evaluating detection and quantitative recovery across clinical backgrounds.

The community experiment complemented this detailed analysis by introducing all 10 taxa simultaneously into all available samples from both cohorts. In addition to providing a more scalable multi-taxon evaluation, this full-cohort design allowed us to assess whether the broad recovery patterns observed in the individual-species subset were preserved across the complete datasets. The 10 taxa were introduced as an equally weighted mixed community at seven total-community spike fractions: 0.01%, 0.05%, 0.1%, 0.5%, 1%, 5%, and 10%. Because each taxon contributed one tenth of the total community spike, the corresponding effective per-taxon fractions were 0.001%, 0.005%, 0.01%, 0.05%, 0.1%, 0.5%, and 1%. For every spiked metagenome, the corresponding unspiked sample was retained as the matched same-background reference for expected-abundance calculations and spike-status comparisons.

### 4.2 Metagenomic preprocessing

Raw metagenomes were processed with a uniform shotgun-metagenomic preprocessing workflow implemented in MetaShotgunPrep v1.0 (https://github.com/microbiomehdlab/MetaShotgunPrep) to minimize technical variation across cohorts before spike-in generation and taxonomic profiling. Raw FASTQ files were inspected with FastQC, adapter- and quality-trimmed with fastp, and filtered to remove low-quality, low-complexity, and short reads. Reads shorter than 60 bp after trimming were discarded. Host-derived reads were removed by alignment to the human reference genome GRCh38 using Bowtie2 in very-sensitive mode, and only unmapped reads were retained. A second FastQC assessment was performed after preprocessing to verify trimming and host-depletion performance.

For each sample, sequencing depth, read retention after preprocessing, and host-filtering summaries were recorded to assess preprocessing performance. Post-QC read-pair depth, read-pair retention, and raw-versus-post-QC depth for the original background metagenomes are summarized in Supplementary Fig. B1. Only high-quality, non-host reads were used for spike-in construction and downstream taxonomic profiling. Exact software versions, command-line parameters, and quality-control thresholds are provided in Supplementary Table A4 and in the pipeline repository.

### 4.3 Selection of spike-in taxa and reference genomes

Ten CRC-associated candidate taxa were selected to form a literature-curated analytical challenge panel (Table 1). The panel was not intended to define a new diagnostic signature or to rank CRC biomarkers by effect size. Instead, it was designed to test whether taxa with prior biological and clinical relevance to CRC microbiome research could be reliably detected and quantified when present as weak, controlled signals in real stool metagenomic backgrounds.

Candidate taxa were prioritized based on three considerations. First, they have been reported in independent CRC stool metagenomic studies, cross-cohort analyses, or meta-analyses of CRC-associated microbiome signatures [1–5, 23–26]. Second, the panel spans ecological and biological contexts relevant to CRC-associated microbiome shifts, including oral-associated anaerobes, gut-associated anaerobes, and taxa reported in studies of the adenoma–carcinoma sequence. Third, the selected organisms were expected to present different analytical challenges for short-read taxonomic profiling, including species- or subspecies-level resolution, database-dependent taxonomic representation, and potential ambiguity relative to closely related or differently named background organisms. This made the panel suitable for evaluating not only detection sensitivity, but also abundance distortion, missed recovery, and off-target taxonomic effects. Reference genomes and assembly accessions for each selected taxon are provided in Table 1.

### 4.4 Spike-in read simulation and insertion

Spike-in read pools were generated from the selected reference genomes using ART [27] with the Illumina HiSeq 2500 error profile (HS25). ART was run in paired-end mode using the `-p` option, generating paired-end (2 × 100)-bp reads. A common read length of 100 bp was used to approximate the sequencing characteristics of the source datasets while keeping the implanted reads identical across cohorts. Reads were generated using a mean fragment length of 350 bp, a fragment-length standard deviation of 10 bp, and 2000× genome coverage. For each target genome, a high-coverage read pool was generated once and subsequently subsampled to obtain the required spike-in fractions. The simulated coverage therefore determined the size of the available source pool and not the final implanted abundance.

The simulated read pools were processed using the same `fastp` quality-trimming, low-complexity-filtering, and minimum-length settings applied to the empirical metagenomes. Quality trimming could shorten reads from their initial length of 100 bp, and reads shorter than 60 bp were removed. Bracken species-level abundance estimation was performed using `-l S, -r 100`, and `-t 16` for all original and spiked metagenomes. Thus, the same preprocessing procedure and fixed 100-bp Bracken read-length model were applied consistently across cohorts and spike-in conditions.

Spike-ins were inserted into the preprocessed, host-depleted empirical metagenomes as paired-end read pairs. ART was run in paired-end mode, and the two reads originating from each simulated fragment were retained together during filtering, subsampling, and insertion. For each empirical sample, *R* was defined as the number of complete background read pairs remaining after preprocessing. For a desired final spike fraction *f*, the number of simulated read pairs added to the sample was

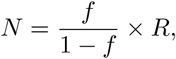

so that the implanted read pairs represented fraction *f* of the final merged paired-end read set.

Individual-species spike-ins were generated at six final read-pair fractions: 0.01%, 0.05%, 0.1%, 0.5%, 1%, and 5%. Community spike-ins were generated at seven total final read-pair fractions: 0.01%, 0.05%, 0.1%, 0.5%, 1%, 5%, and 10%.

In the individual-species experiment, all *N* spike read pairs were drawn from a single taxon-specific pool. In the community experiment, *N* read pairs were distributed equally across the 10 target taxa, giving each taxon an effective implanted fraction of *f/*10. The seven total-community fractions therefore corresponded to effective per-taxon fractions of 0.001%, 0.005%, 0.01%, 0.05%, 0.1%, 0.5%, and 1%.

Simulated spike-in read pairs were merged with the preprocessed empirical metagenomes while leaving the original empirical reads unchanged. Resulting FASTQ files retained metadata identifying the cohort, sample, diagnostic background, spike-in design, target taxon or community label, and nominal final spike fraction.

### 4.5 Taxonomic profiling, abundance representation, and target harmonization

All original and spiked metagenomes were profiled using two widely used workflow–database configurations representing contrasting taxonomic profiling strategies. MetaPhlAn 4 was used as the marker-gene-based profiler, whereas Kraken2 followed by Bracken was used as the k-mer-based classifier and species-level abundance estimator. Kraken2 was run against a database constructed from the Unified Human Gastrointestinal Genome collection, version 2.0.2 (UHGG v2.0.2). The MetaPhlAn 4 database release, software versions, database construction details, and principal profiling parameters are provided in Supplementary Table A5.

To characterize reference-database representation of the implanted genomes, each spike-in assembly was compared with the UHGG v2.0.2 species representatives using Mash. The closest representative was recorded together with its UHGG taxonomic annotation, Mash distance, and Mash-based similarity, defined as 100 × (1 − *D*), where *D* is the Mash distance. This derived similarity was used only as an intuitive summary and was not interpreted as directly measured average nucleotide identity. MetaPhlAn 4 SGBs carrying the target species label or its harmonized taxonomic synonym were identified from the vJan25 database taxonomy. The resulting reference-representation audit is provided in Supplementary Table A2.

The comparison was intentionally restricted to these two workflows and was not intended to survey or rank available taxonomic profilers comprehensively. They were selected because they represent two widely used and contrasting approaches to species-level profiling in human shotgun metagenomics and are routinely used to generate features for downstream microbiome association and biomarker analyses. Their substantial methodological differences therefore make these workflows a useful test case for evaluating the framework and for providing initial evidence that the choice of taxonomic profiler can strongly influence biomarker discovery, with important implications for the identification of reproducible biomarkers.

Species-level abundance profiles were independently generated for each pipeline based on the species-level abundance estimates produced by the corresponding workflow, and spike-recovery performance was assessed using the baseline abundance representation specific to each profiler. Species-level features were retained for downstream analyses, but abundance values were not renormalized after excluding unclassified or above-species assignments, thereby preserving the abundance scale reported by each workflow.

The taxonomic profiles produced by MetaPhlAn 4 and Kraken2/Bracken were analysed in their native label spaces and were not globally merged or standardized. For the 10 implanted targets only, workflow-specific species names, synonyms, and strain- or subspecies-level annotations were mapped to common target identifiers (Supplementary Table A1). This target-specific mapping prevented equivalent taxonomic labels from being counted as missed detections while preserving workflow-specific non-target features. Target detection was defined as a reported species-level abundance greater than zero after mapping, and all labels assigned to the implanted targets were excluded from off-target analyses.

### 4.6 Expected post-spike abundances

Expected post-spike abundances were calculated separately for each profiler using the species-level abundance profile obtained from the corresponding unspiked version of the same sample.

Let *p* denote the profiler, *T_p_* the set of species reported in its species-level abundance profile, and *o_i,p_* the reported abundance of taxon *i* in the corresponding unspiked sample. When an implanted target was not detected in the corresponding unspiked profile, its baseline abundance was defined as *o_s,p_* = 0.

For an individual spike of target taxon *s* at a nominal final read-pair fraction *f*, the expected post-spike abundance of the target under an idealized dilution-only model was

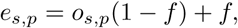

whereas the expected abundance of every non-target taxon (*i* ≠ *s*) was

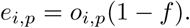

For community spike-ins, the total implanted fraction *f* was divided equally among the 10 target taxa. Each implanted taxon *s* therefore received a taxon-specific fraction

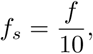

and its expected post-spike abundance was

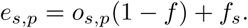

For taxa not included in the implanted community, the expected abundance remained

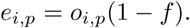

Observed and expected abundances were always compared within the same profiler and relative to the corresponding unspiked sample. Cross-profiler comparisons therefore evaluated recovery relative to each profiler’s native abundance representation rather than assuming equivalence between the complete taxonomic profiles or abundance estimates produced by MetaPhlAn 4 and Kraken2/Bracken.

The dilution-only model represents the idealized expectation that implanted reads are recovered at the intended species level and that the native community changes only through compositional dilution. Deviations from this expectation may reflect incomplete target recovery, abundance-estimation bias, taxonomic misassignment, or profiler-specific compositional effects.

### 4.7 Target recovery metrics

Evaluation focused on metrics relevant to low-abundance biomarker recovery. Metrics were calculated separately for each workflow, cohort, clinical background, spike-in design, target taxon, and spike fraction.

For each target taxon *s* and workflow *p*, detection frequency was defined as the proportion of spiked samples in which the harmonized target abundance was greater than zero. Detection frequencies and recovery-class proportions were reported with Wilson 95% confidence intervals.

Quantitative recovery was evaluated by comparing the observed post-spike abundance (*a_s,p_*) with the expected post-spike abundance (*e_s,p_*) defined in Section 4.6. The observed-to-expected recovery ratio was

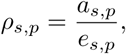

and the absolute relative recovery error was

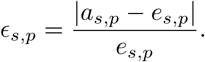

For descriptive recovery-class summaries, observations were classified as *Good* when *ɛ_s,p_* ≤ 0.10, *Intermediate* when 0.10 *< ɛ_s,p_* ≤ 0.50, and *Poor/missed* when *ɛ_s,p_ >* 0.50. When a target was not detected, its observed abundance was set to *a_s,p_* = 0, corresponding to *ρ_s,p_* = 0, *ɛ_s,p_* = 1, and classification as *Poor/missed*. These categories were used for the recovery-class summaries presented in Section 2.2 and Fig. 3C.

Recovery bias was summarized using the median observed-to-expected ratio, whereas recovery variability was summarized using the interquartile range of the observed-to-expected ratios. Absolute relative recovery errors were summarized using medians, interquartile ranges, and their empirical distributions. The same definitions were applied to individual and community spike-ins, with *e_s,p_* calculated using the corresponding individual or effective per-taxon community spike fraction.

### 4.8 Non-target abundance artefacts

To quantify collateral changes in the reported abundance of the native microbial community, all species reported by each profiler except the 10 implanted target taxa (and their harmonized aliases) were treated as non-target taxa. Their observed abundances were compared with the corresponding dilution-only expectations defined in Section 4.6. Let *a_i,p_* denote the observed post-spike abundance of non-target taxon *i* reported by profiler *p*, and *e_i,p_* its expected post-spike abundance, both expressed on the native abundance scale of profiler *p*. For taxa with *e_i,p_ >* 0, the signed relative deviation was defined as

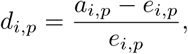

with absolute deviation given by |*d_i,p_*|. Positive values indicated abundance above the dilution-only expectation, whereas negative values indicated abundance below expectation.

Taxa with *e_i,p_* = 0 required separate treatment because their relative deviation was undefined. When *e_i,p_* = 0 and *a_i,p_ >* 0, the taxon was classified as a de novo off-target detection. These observations were excluded only from calculations involving *d_i,p_*, including the mean absolute relative deviation and the standard deviation of signed relative deviation. They were not removed from the profiler-derived abundance tables and were retained as a separate category of putative artefactual appearance for downstream off-target summaries and comparison with differential-abundance results. Taxa with *e_i,p_* = 0 and *a_i,p_* = 0 were absent from both the expected and observed profiles and contributed neither to deviation summaries nor to de novo-detection counts.

For each profiler, spike fraction, and non-target taxon with *e_i,p_ >* 0, abundance distortion across matched samples was summarized using the mean absolute relative deviation,

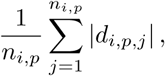

and the standard deviation of the signed relative deviations,

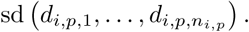

A non-target taxon was operationally classified as artefact-prone when either its mean absolute relative deviation exceeded 5% or the standard deviation of its signed relative deviation exceeded 5%. De novo off-target detections were retained as a separate artefact category rather than being assigned an arbitrary relative-deviation value.

### 4.9 Differential-abundance recovery and biomarker metrics

The propagation of profiler behaviour into downstream biomarker discovery was evaluated using an intentionally unpaired group-level differential-abundance analysis. This design was selected to reflect the inferential objective of conventional microbiome biomarker-discovery studies, in which a candidate taxon must produce a sufficiently consistent abundance difference across groups to emerge as a biomarker. The objective was therefore not to test whether the implanted signal increased within every individual sample, but whether the workflow recovered the signal consistently enough for it to become statistically detectable at the group level.

For each profiler, cohort, diagnostic background, spike-in design, and spike fraction, the set of spiked abundance profiles was compared with the corresponding set of unspiked profiles using MaAsLin2. Spike status was included as the fixed effect of interest, with the unspiked state as the reference category. Although each spiked profile originated from the same biological background as a corresponding unspiked profile, biological-sample identity was not included as a random effect in the primary analysis because the intended endpoint was group-level biomarker recoverability rather than estimation of a within-sample spike effect.

Differential-abundance models were fitted using MaAsLin2 version 1.18.0 under R version 4.3.3, with linear-model analysis, no additional normalization, logarithmic transformation, Benjamini–Hochberg correction, and a minimum prevalence threshold of 0.10 within each fitted model. No external abundance-, prevalence-, or variance-based feature filtering was applied before MaAsLin2 in the original analysis mode; all profiler-reported species-level taxon columns were passed to MaAsLin2.

For individual spike-ins, the implanted taxon was the only true-positive feature. For community spike-ins, the true-positive set comprised the 10 implanted community members after target-specific alias harmonization. All taxa outside the implanted target set were treated as non-target features.

An implanted target was considered recovered as a differential-abundance biomarker when it was significantly enriched in the spiked profiles relative to the matched unspiked group. Significance was defined as a false-discovery-rate-adjusted *q*-value of *q* ≤ 0.10, together with a positive spike-status coefficient.

The minimum recoverable spike fraction was defined as the lowest tested fraction at which the implanted target satisfied both the significance and coefficient-direction criteria. Targets that were not significantly enriched at any tested fraction were classified as not recovered.

Off-target burden was defined as the number of non-target taxa that were significantly enriched in the spiked profiles relative to the matched unspiked group. Significant non-target associations with negative coefficients were not counted as false-positive enriched biomarkers and, where relevant, were summarized separately as spike-associated depletions. In community-spike analyses, significant profiler-specific labels mapping to the same implanted taxon were collapsed to the corresponding canonical target before target recovery was summarized.

For the supplementary taxon-specific off-target analysis, recurrence was defined as the fraction of eligible differential-abundance contexts in which a non-target feature was significantly enriched, calculated separately by profiler, spike-in design, implanted target, and spike fraction. Eligible contexts comprised the available cohort and diagnostic-background combinations. For individual-species spike-ins, the maximum recurrence across tested spike fractions was used to summarize each implanted-target–off-target feature pair. Same-genus representation among enriched off-target feature-context observations was compared with that among non-enriched non-target observations using descriptive odds ratios with a 0.5 continuity correction and approximate 95% confidence intervals. For community spike-ins, off-target recurrence was joined to the corresponding dilution-model abundance-error summaries. Intersections between individual-species and community designs included features recurring in at least 20% of their eligible contexts.

Biomarker-recovery precision, recall, and F1 score were calculated as

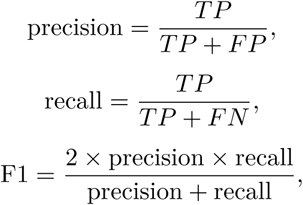

where *TP* denotes the number of implanted target taxa recovered as significantly enriched biomarkers, *FP* denotes the number of significantly enriched non-target taxa, and *FN* denotes the number of implanted targets that were not recovered. For community spike-ins, *TP* + *FN* = 10. When no taxa were called significant and the denominator of the precision or F1 calculation was zero, the corresponding metric was defined as zero.

### 4.10 Recovery-related drivers of biomarker significance

Associations between differential-abundance significance and recovery-related variables were evaluated at the weakest individual spike fraction, 0.01%. For each implanted target, biomarker significance was represented as (− log_10_(*q*)), where *q* was the false-discovery-rate-adjusted value obtained from the spike-status MaAsLin2 analysis comparing spiked and unspiked profiles.

The evaluated recovery-related variables were the target abundance in the corresponding unspiked profile, target detection frequency, median absolute relative recovery error, and recovery variability, defined as the interquartile range of the observed-to-expected recovery ratios. Spearman rank correlations were calculated separately by profiler, cohort, and clinical background. Continuous variables were additionally divided into tertiles for visualization only; all correlation analyses used the original continuous values.

### 4.11 Artefact-aware exclusion of off-target differential-abundance calls

Artefact-aware exclusion was evaluated using the community spike-in experiment. Again, because only the 10 implanted community members should increase after spike insertion, all remaining taxa were treated as non-target taxa and were expected to change only according to the dilution-only expectation defined in Section 4.8.

For each profiler, spike fraction, and non-target taxon, abundance distortion was summarized from the sample-level relative deviations. The two summary statistics used for artefact classification were the mean absolute relative deviation and the standard deviation of the signed relative deviation. At least 20 finite sample-level relative-error observations were required for a profiler–fraction–taxon estimate to be eligible for artefact classification. A taxon was classified as artefact-prone at threshold *τ* when

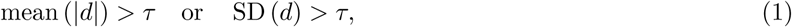

where *d* denotes the signed relative deviation from the dilution-only expectation defined in Section 4.8. The primary analysis used *τ* = 0.05, corresponding to a 5% relative-error threshold. Sensitivity analyses were additionally performed using thresholds of 2.5%, 10%, and 20%. Artefact classification was performed separately for each profiler and spike fraction; a taxon was not required to exceed the threshold at multiple spike fractions.

The implanted community targets and all profiler-specific labels mapped to those targets were designated as target features before exclusion and were therefore protected from removal. Consequently, artefact-aware exclusion acted only on enriched non-target differential-abundance calls and could not remove a canonical implanted target or one of its mapped aliases by construction.

Two complementary evaluations were performed. For the internal analysis, artefact statistics were pooled across both cohorts and across diagnostic backgrounds, while remaining specific to profiler, spike fraction, and taxon. The resulting pooled profiler- and fraction-specific artefact lists were applied to differential-abundance results from the same overall community-spike evaluation.

For the reciprocal cross-study transfer analysis, artefact statistics were calculated separately within each cohort, pooling diagnostic backgrounds within that cohort. Profiler- and fraction-specific artefact lists learned in one cohort were then frozen and applied without modification to differential-abundance calls in the other cohort. Transfer was evaluated in both directions between the FengQ 2015 and ZellerG 2014 cohorts.

In both evaluations, exclusion was applied post hoc to the previously identified differential-abundance results generated from the original, unfiltered feature tables. MaAsLin2 models were not refitted after feature exclusion. Eligible differential-abundance calls were restricted to features with a positive spike-status coefficient and a false-discovery-rate-adjusted value of *q* ≤ 0.10, as defined in Section 4.9. Each evaluation context comprised one profiler, study, diagnostic background, and spike fraction.

For each context, off-target burden was defined as the number of significantly enriched non-target taxa remaining before or after artefact-aware exclusion. Differential-abundance-list precision was calculated using the definition in Section 4.9, where *TP* denotes the number of recovered canonical implanted targets and *FP* denotes the number of significantly enriched non-target taxa remaining after optional artefact-aware exclusion. F1 score was calculated using the same precision definition together with the canonical-target recall defined in Section 4.9. Because implanted targets and their aliases were protected from exclusion, target recall was unchanged by construction. The filtering analysis was therefore evaluated primarily through changes in off-target burden, precision, and F1 score.

### 4.12 Evaluation of spike-informed abundance calibratability

We evaluated whether the relationship between expected and profiler-reported target abundance was sufficiently structured to support future spike-informed abundance calibration. Analyses used the community spike-in experiment and were performed separately for each profiler–target combination, pooling observations across cohorts, diagnostic backgrounds, and community spike fractions for the primary calibratability assessment.

Expected total post-spike abundance was calculated as described in Section 4.6. For biological sample *i*, community target taxon *s*, profiler *p*, and total community spike fraction *f_i_*, the expected abundance was

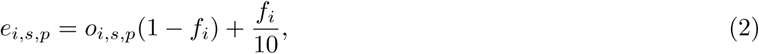

where *o_i,s,p_* denotes the abundance reported by profiler *p* for taxon *s* in the matched unspiked profile. The term *f_i_/*10 reflects the equal allocation of the implanted community fraction among the 10 target taxa. The observed abundance, *a_i,s,p_*, was the species-level reported abundance reported for the corresponding target in the spiked profile.

Expected and observed abundances were transformed as

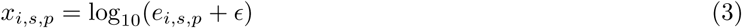

and

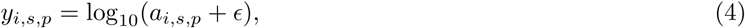

where *ɛ* was defined as one half of the smallest positive expected or observed abundance in the calibratability-analysis dataset. The logarithmic transformation was used because target abundances spanned several orders of magnitude. For visualization, fitted values were back-transformed and displayed on the original percentage-abundance scale.

Target detection was summarized separately from quantitative abundance-response modelling. Quantitative models included only observations for which the target was detected, and both expected and observed abundances were positive. Non-detection was therefore not treated as a calibratable positive-abundance measurement.

For visualization of the forward abundance-response relationship, generalized additive models were fitted as

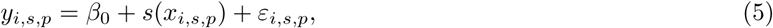

where *s*(·) denotes a penalized cubic regression spline. Models were fitted using the mgcv package with restricted maximum-likelihood estimation. The maximum spline basis dimension was *k* = 5, with a smaller basis used when the number of distinct predictor values was insufficient to support this value.

Predictive calibratability was evaluated by comparing three approaches for estimating expected abundance from profiler-reported abundance. The first was a raw identity prediction, in which the observed abundance was used directly as the prediction of expected abundance. The second was an ordinary linear regression fitted on the logarithmic scale,

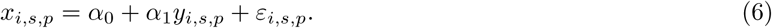

The third was an inverse generalized additive model,

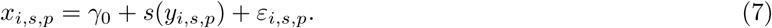

Predictive performance was assessed using five-fold grouped cross-validation repeated five times with random seed 1. The original biological-sample identifier was used as the grouping unit, ensuring that all observations derived from the same stool sample remained within the same fold. A profiler–taxon combination was eligible for quantitative modelling when the target had an overall detection rate of at least 0.50, at least 40 detected observations, at least 80 positive-abundance observations, and positive observations from at least eight original biological samples.

Held-out performance was calculated separately for each fold and repeat and then summarized as the median across cross-validation partitions. Performance metrics included median absolute log-scale error,

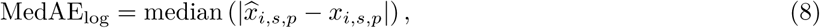

root-mean-square log-scale error,

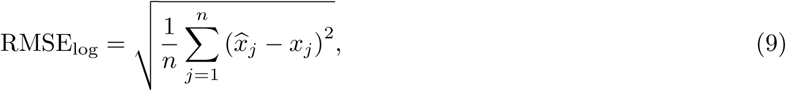

median absolute relative error, Spearman rank correlation, and the proportions of predictions falling within 10% and 50% of the expected abundance. Improvement was defined as the reduction in held-out median absolute log-scale error relative to the raw identity prediction,

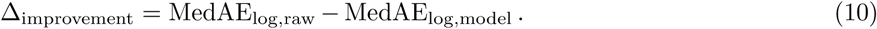

Positive values of Δ_improvement_ therefore indicated that the fitted calibration model reduced held-out prediction error relative to using the profiler-reported abundance without calibration.

Profiler–taxon combinations were assigned to four descriptive calibratability categories using the held-out performance of the inverse GAM. A combination was classified as not assessable (ND) when the overall detection rate was below 0.50, fewer than 40 detected observations were available, fewer than 80 positive-abundance observations were available, fewer than eight biological samples contributed positive observations, or no finite grouped cross-validation estimate could be obtained. Among assessable combinations, strong evidence of calibratability (S) required an improvement in median absolute log-scale error greater than 0.10 together with a residual GAM median absolute log-scale error below 0.25. Conditional evidence (C) was assigned when the GAM produced a positive improvement but did not satisfy both strong-evidence criteria. Little evidence (L) was assigned when the GAM did not reduce held-out median absolute log-scale error relative to the raw identity prediction. These categories were used as descriptive summaries of calibration benefit and were not interpreted as universally validated calibration thresholds.

Cross-study transferability was evaluated by fitting the inverse GAM in one cohort and applying it without refitting to the other cohort. This analysis was conducted in both directions between the FengQ 2015 and ZellerG 2014 cohorts. Transfer models required at least 30 positive-abundance observations in the training cohort and at least 10 in the test cohort, and performance was evaluated using the same log-scale prediction-error metrics.

The purpose of this analysis was to determine whether profiler-specific abundance distortion was reproducible and potentially calibratable. The fitted models were used to evaluate abundance-response predictability and cross-cohort transferability rather than to transform the original unspiked cohort profiles.

### 4.13 Computational environment and reproducibility

Computational workflows were executed on the LOBO high-performance computing cluster under the Slurm workload manager. Compute nodes were equipped with two Intel Xeon E5-2630 v4 processors, providing 20 physical cores and 40 hardware threads per node, and either 126 or 254 GB of RAM, depending on the node class. Taxonomic profiling, spike-in generation, and command-line processing were performed in Apptainer containers using version-controlled micromamba or conda software environments. Statistical analyses and figure generation were performed using R version 4.3.3.

Software and package versions, reference-database releases, container identifiers, command-line parameters, analysis settings, and random seeds are documented in the supplementary tables and in the project repository.

Source code, workflow scripts, software-environment specifications, spike-in metadata and target-alias mappings are available at https://github.com/microbiomehdlab/ground-truth-taxonomic-calibration.

## Ethics and data availability

All human metagenomic datasets analyzed in this study were obtained from publicly accessible repositories and were reprocessed in accordance with the original informed consent provisions and data-sharing agreements. Accession identifiers and cohort-level metadata are provided in the project repository.

## Contributions

A.S., A.T.F. and A.S.A. conceived and designed the study. A.S. developed the computational pipeline and performed the analyses. C.R.T. reproduced the analyses across spike-in conditions and datasets. A.S. drafted the manuscript, and all authors contributed to the interpretation of the results, critically revised the manuscript, and approved the final version.

## Acknowledgements

This work was supported by GIMM-CARE, which is funded by the European Union’s Horizon Europe research and innovation programme under grant agreement No. 101060102. GIMM-CARE is co-funded by the Portuguese Government; the Fundação para a Ciência e a Tecnologia (FCT); the Francisco Manuel dos Santos Society Group (ARICA–Investimentos, Participações e Gestão and Jeŕonimo Martins); GIMM; and CAML–Lisbon Academic Medical Centre (https://doi.org/10.3030/101060102). Additional support was provided by national funds through FCT under the Associate Laboratory programme (LA/P/0082/2020; https://doi.org/10.54499/LA/P/0082/2020) and the R&D Unit funding programme (UID/06357/2025; https://doi.org/10.54499/UID/06357/2025). AS is supported by FCT grant 2025.02293.BD, and ASA by FCT grant 2021.02791.CEECIND. This work was also supported by INESC-ID and national funds through FCT under projects UID/50021/2025 (https://doi.org/10.54499/UID/50021/2025) and UID/PRR/50021/2025 (https://doi.org/10.54499/UID/PRR/50021/2025).

The authors gratefully acknowledge the support & assistance of the Advanced Data Analysis facility at GIMM, especially A. Barros. Generative AI tools, specifically ChatGPT and Codex (OpenAI), were used to assist with code development, debugging, and analytical workflows, as well as with manuscript preparation and language refinement. The authors retain full responsibility for the scientific content, analyses, and final manuscript. All analytical outputs were validated by the authors.

