## Supplementary Information for "Ground-truth *in silico* spike-ins reveal limits of microbiome biomarker recovery in colorectal cancer"

### Appendix A Supplementary Methods and Tables

**Table A1**: **Complete profiler-specific taxonomic-label harmonization used for spike-in target analyses.** The canonical target corresponds to the taxonomic designation recorded in the spike-in design. Identity mappings are shown explicitly for completeness. The *Fusobacterium nucleatum* spike-in used a subspecies-annotated reference assembly, whereas both profilers reported the target at species level.

| **Canonical target** | **MetaPhlAn 4 label** | **Kraken2/Bracken label** |
|---|---|---|
| *Bacteroides fragilis* | *Bacteroides fragilis* | *Bacteroides fragilis* |
| *Clostridium symbiosum* | *Clostridium symbiosum* | `Clostridium_Q symbiosum` |
| *Dialister pneumosintes* | *Dialister pneumosintes* | *Allisonella pneumosintes* |
| *Fusobacterium nucleatum* subsp. *nucleatum* | *Fusobacterium nucleatum* | *Fusobacterium nucleatum* |
| *Hungatella hathewayi* | *Hungatella hathewayi* | `Hungatella_A hathewayi_A` |
| *Parvimonas micra* | *Parvimonas micra* | *Parvimonas micra* |
| *Peptostreptococcus anaerobius* | *Peptostreptococcus anaerobius* | *Peptostreptococcus anaerobius* |
| *Peptostreptococcus stomatis* | *Peptostreptococcus stomatis* | *Peptostreptococcus stomatis* |
| *Porphyromonas asaccharolytica* | *Porphyromonas asaccharolytica* | *Porphyromonas asaccharolytica* |
| *Prevotella intermedia* | *Prevotella intermedia* | *Prevotella intermedia* |

**Table A2**: **Representation of implanted target genomes in the profiler reference databases.** For each target genome, the closest UHGG v2.0.2 species representative was identified using Mash. Mash-based similarity was calculated as $100 \times (1 - D)$, where $D$ is the Mash distance, and is provided as an intuitive summary rather than as a direct measurement of average nucleotide identity. MetaPhlAn 4 SGBs were identified by matching the target species or its harmonized taxonomic synonym to the vJan25 database taxonomy. Multiple SGBs indicate that multiple SGBs carried the corresponding species-level label and do not represent multiple sequence-based assignments of the spike-in assembly.

| **Target** | **Spike-in assembly** | **Closest UHGG representative** | **UHGG species label** | **Mash distance** | **Mash-based similarity** | **MetaPhlAn vJan25 SGB(s)** |
|---|---|---|---|---|---|---|
| Bfrag | GCF_000025985.1 | MGYG000001337 | *Bacteroides fragilis* | 0.0128 | 98.72% | SGB1853; SGB1855; SGB104919 |
| Csym | GCF_000466485.1 | MGYG000001367 | *Clostridium Q symbiosum* | 0.0126 | 98.74% | SGB4699; SGB161423 |
| Dpne | GCF_003570845.1 | MGYG000001613 | *Allisonella pneumosintes* | 0.0050 | 99.50% | SGB5842 |
| Fnuc | GCF_003019295.1 | MGYG000001459 | *Fusobacterium nucleatum* | 0.0102 | 98.98% | SGB6011; SGB6013 |
| Hhat | GCF_000235505.1 | MGYG000001688 | *Hungatella A hathewayi A* | 0.0000 | 100.00% | SGB4739; SGB4741; SGB4742 |
| Pmic | GCF_003454775.1 | MGYG000001301 | *Parvimonas micra* | 0.0207 | 97.93% | SGB6649; SGB6653 |
| Pana | GCF_900454605.1 | MGYG000000296 | *Peptostreptococcus anaerobius* | 0.0118 | 98.82% | SGB746 |
| Psto | GCA_000147675.2 | MGYG000004565 | *Peptostreptococcus stomatis* | 0.0164 | 98.36% | SGB748 |
| Porp | GCF_000183605.1 | MGYG000004267 | *Porphyromonas asaccharolytica* | 0.0174 | 98.26% | SGB1981 |
| Pint | GCF_002797175.1 | MGYG000004629 | *Prevotella intermedia* | 0.0345 | 96.55% | SGB1560 |

**Table A3**: Samples included in the individual-species spike-in experiment. The table lists the original cohort identifiers and corresponding ENA run accessions for the balanced subset of 10 samples per clinical group from each cohort. Rows are grouped for presentation and do not imply the order in which samples were selected.

| Cohort and clinical group | Original sample ID | ENA run accession |
|---|---|---|
| **FengQ 2015 — Control** | | |
| | SID31700 | ERR688559 |
| | SID31711 | ERR688561 |
| | SID31714 | ERR688562 |
| | SID31723 | ERR688563 |
| | SID31749 | ERR688564 |
| | SID31750 | ERR688565 |
| | SID31766 | ERR688566 |
| | SID31300 | ERR688528 |
| | SID31328 | ERR688529 |
| | SID31333 | ERR688530 |
| **FengQ 2015 — Adenoma** | | |
| | SID31705 | ERR688560 |
| | SID530002 | ERR688587 |
| | SID530018 | ERR688584 |
| | SID530026 | ERR688585 |
| | SID530028 | ERR688586 |
| | SID530039 | ERR688588 |
| | SID31030 | ERR688508 |
| | SID31137 | ERR688512 |
| | SID31455 | ERR688545 |
| | SID31477 | ERR688546 |
| **FengQ 2015 — CRC** | | |
| | SID31873 | ERR688574 |
| | SID31874 | ERR688575 |
| | SID31875 | ERR688576 |
| | SID31876 | ERR688577 |
| | SID31877 | ERR688578 |
| | SID31878 | ERR688579 |
| | SID31879 | ERR688580 |
| | SID31880 | ERR688581 |
| | SID31881 | ERR688582 |
| | SID31883 | ERR688583 |
| **ZellerG 2014 — Control** | | |
| | CCIS00146684ST-4-0 | ERR478961 |
| | CCIS00281083ST-3-0 | ERR478963 |
| | CCIS02124300ST-4-0 | ERR478967 |
| | CCIS02856720ST-4-0 | ERR478973 |
| | CCIS03473770ST-4-0 | ERR478975 |
| | CCIS03857607ST-4-0 | ERR478979 |
| | CCIS05314658ST-4-0 | ERR478983 |
| | CCIS07277498ST-4-0 | ERR478989 |
| | CCIS07539127ST-4-0 | ERR478993 |
| | CCIS07648107ST-4-0 | ERR478997 |
| **ZellerG 2014 — Adenoma** | | |
| | CCIS08668806ST-3-0 | ERR478999 |
| | CCIS10793554ST-4-0 | ERR479009 |
| | CCIS11019776ST-4-0 | ERR479013 |
| | CCIS19142497ST-3-0 | ERR479058 |
| | CCIS22275061ST-4-0 | ERR479070 |

Continued on next page

Table A3 – continued from previous page

| Cohort and clinical group | Original sample ID | ENA run accession |
|---|---|---|
| | CCIS22906510ST-20-0 | ERR479076 |
| | CCIS24898163ST-4-0 | ERR479087 |
| | CCIS25399172ST-4-0 | ERR479090 |
| | CCIS28384594ST-4-0 | ERR479098 |
| | CCIS31434951ST-20-0 | ERR479104 |
| **ZellerG 2014 — CRC** | | |
| | CCIS02379307ST-4-0 | ERR478969 |
| | CCIS06260551ST-3-0 | ERR478985 |
| | CCIS11015875ST-4-0 | ERR479011 |
| | CCIS12370844ST-4-0 | ERR479028 |
| | CCIS12656533ST-4-0 | ERR479032 |
| | CCIS14449628ST-4-0 | ERR479038 |
| | CCIS15704761ST-4-0 | ERR479040 |
| | CCIS17669415ST-4-0 | ERR479056 |
| | CCIS21278152ST-4-0 | ERR479066 |
| | CCIS22416007ST-4-0 | ERR479074 |

**Table A4: fastp trimming parameters used in the MetaShotgunPrep preprocessing workflow.**

| Parameter | Purpose | Rationale |
|---|---|---|
| `--detect_adapter_for_pe` | Automatically detects adapters in paired-end reads | Allows adapter removal in mixed sequencing batches where library-specific adapter sequences may not be known a priori. |
| `--cut_front --cut_tail` | Removes low-quality bases from both read ends | Reduces terminal sequencing errors, which are common in Illumina reads and can affect downstream mapping and taxonomic classification. |
| `--cut_window_size 4` | Performs sliding-window quality trimming | Provides local quality trimming while avoiding excessive read shortening. |
| `--cut_mean_quality 20` | Requires a minimum mean quality score of Q20 within the trimming window | Q20 corresponds to an approximate base-call error rate of 1%, providing a conservative threshold for metagenomic preprocessing. |
| `--trim_poly_g` | Removes poly-G artefacts | Reduces NovaSeq/NextSeq poly-G tail artefacts that may otherwise introduce spurious sequence signal. |
| `--low_complexity_filter` | Removes low-complexity reads | Filters repetitive or uninformative reads that can contribute to false-positive taxonomic assignments. |
| `--complexity_threshold 30` | Sets the minimum sequence-complexity threshold | Removes low-complexity artefacts while retaining most biologically informative reads. |
| `--length_required 60` | Discards reads shorter than 60 bp after trimming | Excludes very short reads, which map less reliably and may inflate false-positive taxonomic classifications. |

**Table A5**: **Taxonomic profiling tools, databases, and key settings used for species-level abundance estimation.** All profilers were run inside the `taxonomic-tools` container to ensure software reproducibility. Databases were stored outside the container and bind-mounted at runtime.

| Component | Software / database | Version or release | Use in this study |
|---|---|---|---|
| Container environment | - | Docker image tag `0.6`; conda environment `taxonomic_tools` | Provided a fixed execution environment containing `kraken2`, `bracken`, `metaphlan`, `bowtie2`, `fastqc`, `fastp`. The container was run with Docker or Apptainer/Singularity depending on the compute environment. |
| MetaPhlAn | MetaPhlAn 4 | MetaPhlAn 4.2.2 with vJan25 ChocoPhlAnSGB database release `ChocoPhlAnSGB_202503` | Used as the marker-based taxonomic profiler. Species-level reported abundance profiles were generated from clade-specific marker genes. |
| MetaPhlAn database | ChocoPhlAnSGB | vJan25 / `ChocoPhlAnSGB_202503` | Downloaded once into an external database directory using `metaphlan --install --db_dir`. The database directory was bind-mounted into the container at runtime. |
| Kraken2 | Kraken2 | Version 2.1.6 | Used as the k-mer-based read classifier. Reads were classified against the UHGG Kraken2 database, and Kraken2 reports were used as input for Bracken. |
| Kraken2 database | Unified Human Gastrointestinal Genome database | UHGG v2.0.2 Kraken2 database | Used as the gut-specific Kraken2 reference database. The database was downloaded from the EBI/MGnify FTP release and stored externally to the container. |
| Bracken | Bracken | Version 3.0.1 | Used to re-estimate species-level abundances from Kraken2 classification reports. Species-level estimation was performed with `-l S`, a read length of `-r 100`, and a minimum-read threshold of `-t 16`. |
| Bracken k-mer distributions | UHGG v2.0.2 Bracken distributions | Prebuilt distributions included with the UHGG v2.0.2 Kraken2 database; distributions available for 50, 100, 150, 200, and 250 bp | Species-level abundance estimation was performed with `-l S`. The read-length parameter `-r` was set to match the processed read length used for Bracken abundance estimation. |
| ART | ART | Version 2.5.8 | Used to simulate paired-end spike-in read pools with the Illumina HiSeq 2500 error profile. |
| fastp | fastp | Version 0.23.4 | Used for adapter removal, quality trimming, low-complexity filtering, and minimum-length filtering. |
| Bowtie2 | Bowtie2 | Version 2.5.4 | Used for host-read removal. |
| FastQC | FastQC | Version 0.12.1 | Used for pre- and post-processing read-quality assessment. |
| Container runtime | Apptainer | version 1.1.3 | Container execution on LOBO. |

**Table A6**: **MaAsLin2 configuration used for the original unpaired spike-in differential-abundance analysis.**

| Component | Setting | Value or implementation |
|---|---|---|
| Input feature level | Taxonomic rank | Species-level profiler abundances. |
| Input abundance scale | Feature values | Relative-abundance proportions after profiler-specific post-processing. |
| Normalization | `normalization` | `NONE`; no additional normalization was applied within MaAsLin2. |
| Transformation | `transform` | `LOG`. |
| Statistical method | `analysis_method` | `LM`. |
| Minimum abundance | `min_abundance` | 0. |
| Minimum prevalence | `min_prevalence` | 0.10. |
| Minimum variance | `min_variance` | 0. |
| Predictor standardization | `standardize` | `TRUE`. |
| Fixed effect | `fixed_effects` | Spike status. |
| Reference category | `reference` | Unspiked sample. |
| Random effect | `random_effects` | None. |
| Model structure | Unpaired linear model | Spike status was fitted as the fixed effect of interest; no biological-sample random intercept was included. |
| Analysis stratification | Separate model sets | Models were fitted separately for each profiler, study, diagnostic background, spike design, and spike fraction. |
| Coefficient direction | Enrichment criterion | Positive spike-status coefficient. |
| Multiple-testing method | `correction` | Benjamini–Hochberg correction. |
| Significance threshold | Adjusted significance | $q \leq 0.10$. |
| Correction scope | Hypothesis family | Multiple-testing correction was performed separately within each fitted model. |
| Zero handling | Zero-valued abundances | No external zero replacement or pseudocount addition was applied before MaAsLin2; zero handling followed the package's built-in `LOG`-transformation procedure. |
| Pre-MaAsLin2 feature filtering | External filtering | No external abundance-, prevalence-, or variance-based feature filtering was applied in the original analysis mode; all profiler-reported taxon columns were passed to MaAsLin2. |
| Internal feature filtering | MaAsLin2 filtering | Features were evaluated using `min_abundance = 0`, `min_prevalence = 0.10`, and `min_variance = 0`. |
| Software | MaAsLin2 version | MaAsLin2 version 1.18.0 under R version 4.3.3. |
| Random seed | Reproducibility | Not applicable to MaAsLin2 linear-model fitting; stochastic procedures were not used. |

**Table A7**: **Study design, sample usage, spike fractions, and analysis scale.**

| Component | Setting | Value or implementation |
|---|---|---|
| Cohorts | Biological backgrounds | FengQ 2015 and ZellerG 2014. |
| Diagnostic backgrounds | Clinical groups | Control, adenoma, and colorectal carcinoma. |
| Profilers | Taxonomic workflows | Kraken2/Bracken and MetaPhlAn 4. |
| Spike-in taxa | Target panel | Ten CRC-associated taxa listed in Table 1. |
| Spike-in read type | Simulated reads | Paired-end $2 \times 100$ bp reads. |
| Spike-in simulator | Read simulation | ART using the Illumina HiSeq 2500 error profile. |
| Spike-in fragment model | Insert-size parameters | Mean fragment length of 350 bp and fragment-length standard deviation of 10 bp. |
| Spike-in pool depth | Simulation depth | $2000\times$ genome coverage per target genome before deterministic subsampling. |
| Individual spike-in design | Sample subset | Ten samples per diagnostic group per cohort. |
| Individual spike-in scale | Original sample backgrounds | 2 cohorts $\times$ 3 diagnostic groups $\times$ 10 samples = 60 original sample backgrounds. |
| Individual spike-in targets | Spike design | Each of the ten target taxa was implanted separately. |
| Individual spike-in fractions | Final read-pair fractions | 0.01%, 0.05%, 0.1%, 0.5%, 1%, and 5%. |
| Individual spike-in profiles | Spiked metagenomes | 60 sample backgrounds $\times$ 10 taxa $\times$ 6 fractions = 3,600 spiked metagenomes. |
| Community spike-in design | Sample subset | All available samples from both cohorts. |
| Community spike-in scale | Original sample backgrounds | $61 + 47 + 46 = 154$ FengQ 2015 samples and $61 + 42 + 53 = 156$ ZellerG 2014 samples; 310 original sample backgrounds in total. |
| Community spike-in targets | Spike design | All ten target taxa were implanted simultaneously as an equal-weight mixed community. |
| Community total fractions | Final read-pair fractions | 0.01%, 0.05%, 0.1%, 0.5%, 1%, 5%, and 10%. |
| Community effective fractions | Effective per-taxon fractions | 0.001%, 0.005%, 0.01%, 0.05%, 0.1%, 0.5%, and 1.0%. |
| Community spike-in profiles | Spiked metagenomes | 310 sample backgrounds $\times$ 7 fractions = 2,170 spiked metagenomes. |
| Total spike-in profiles | Combined spike-in dataset | 3,600 + 2,170 = 5,770 spiked metagenomes. |
| Unspiked references | Same-background controls | Each original metagenome was retained as the corresponding unspiked reference. |
| Expected-abundance model | Ground-truth calculation | Profiler-specific baseline abundance plus the known spike fraction under the dilution-only model. |
| Differential-abundance analysis | Comparison structure | Spiked profiles were compared with unspiked profiles within profiler, study, diagnostic background, spike design, and spike fraction. |

### Appendix B Supplementary Figures

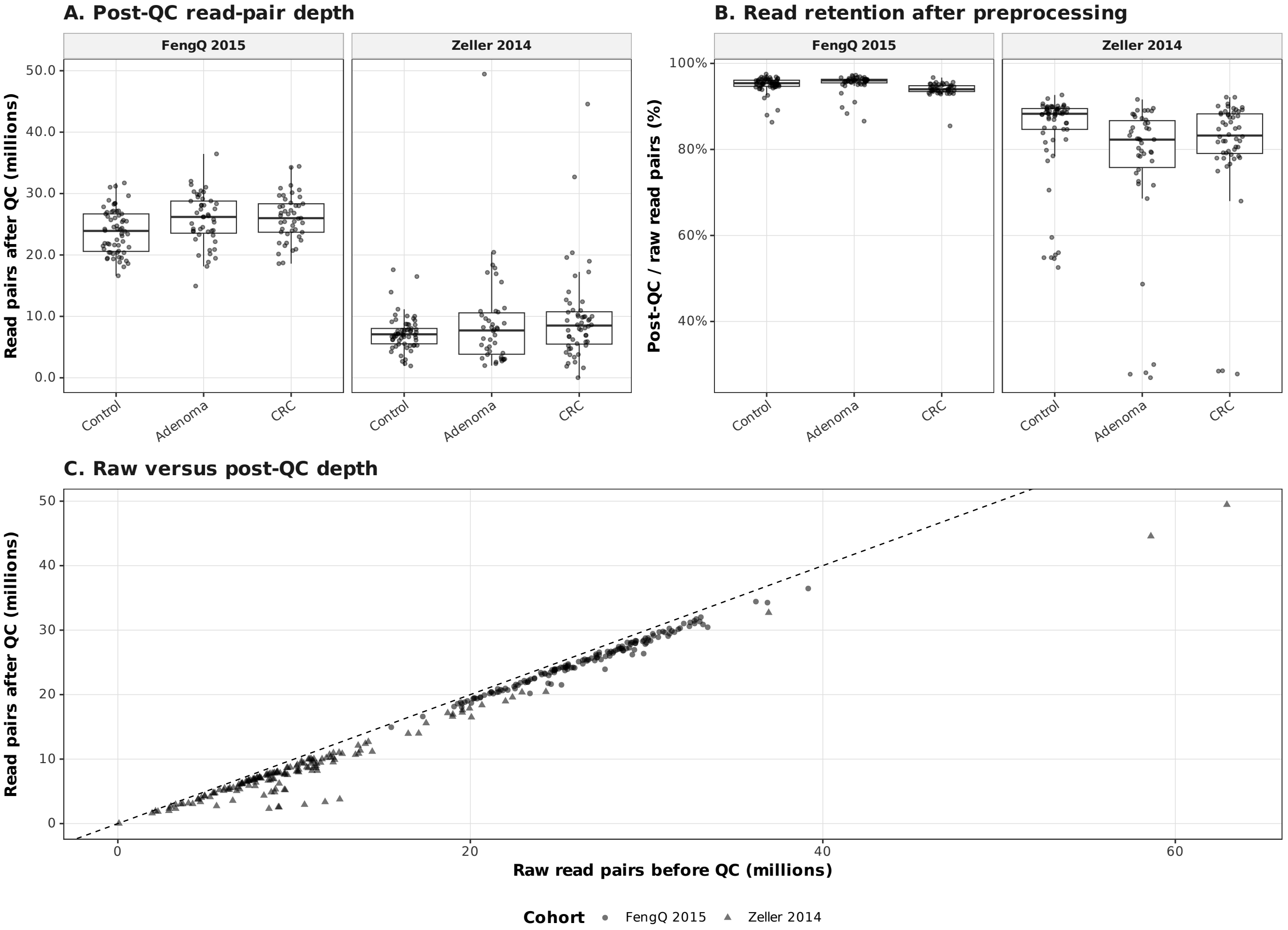


**Fig. B1: Sequencing depth and read retention after preprocessing in the two validation cohorts. (A)** Post-QC paired-end sequencing depth, expressed as millions of read pairs, across diagnostic groups in the FengQ 2015 and ZellerG 2014 cohorts. **(B)** Percentage of raw read pairs retained after preprocessing for the same samples and diagnostic groups. In panels A and B, points represent individual samples and boxplots summarize the corresponding distributions. **(C)** Relationship between raw and post-QC sequencing depth for individual samples. The dashed diagonal denotes equality between raw and post-QC depth. Point shape identifies cohort.

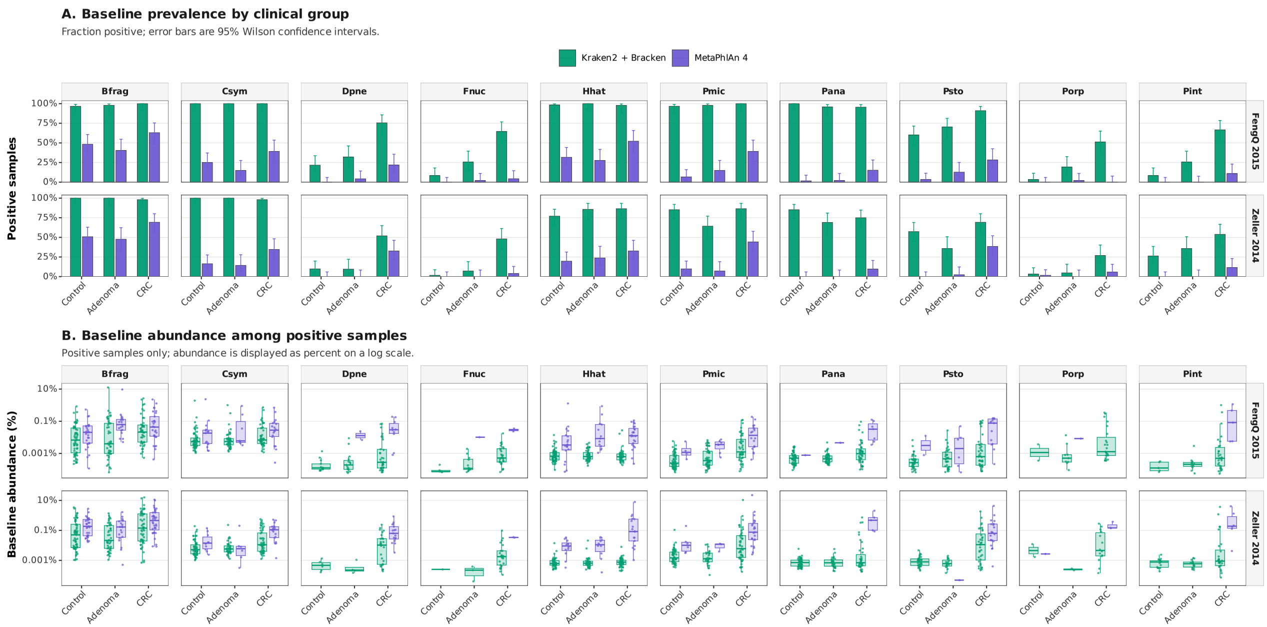


**Fig. B2: Profiler-dependent baseline detection and abundance of all ten CRC-associated target taxa. (A)** Full-cohort baseline prevalence of each target taxon across diagnostic groups in the FengQ 2015 and ZellerG 2014 cohorts, stratified by profiler. Bars show the fraction of samples with non-zero baseline abundance, and error bars indicate 95% Wilson confidence intervals. **(B)** Baseline abundance distributions among positive samples, defined separately for each profiler–taxon combination as samples with abundance greater than zero. Boxplots summarize the distributions and points represent individual samples. Abundances are displayed as percentages on a logarithmic scale. Taxa are ordered consistently with the main figures.

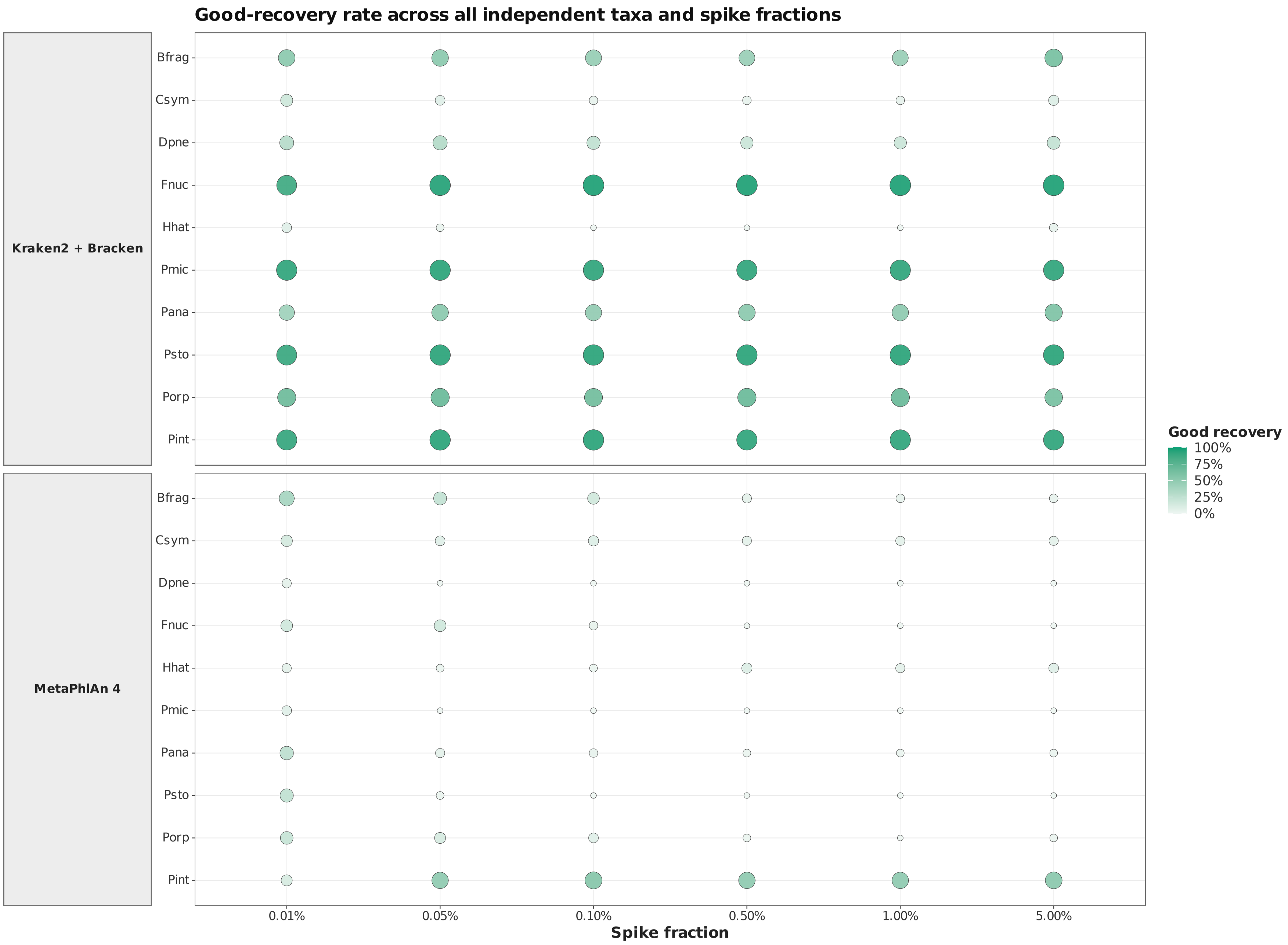


**Fig. B3: Good quantitative recovery of independently spiked target taxa across all tested spike fractions.** The percentage of samples exhibiting Good quantitative recovery is shown for each target taxon, profiler, and independent spike fraction. Good recovery was defined as an absolute relative recovery error $\leq 10\%$, where relative recovery error was calculated from the observed post-spike abundance relative to the expected post-spike abundance. Both symbol size and colour intensity encode the percentage of samples satisfying this criterion.

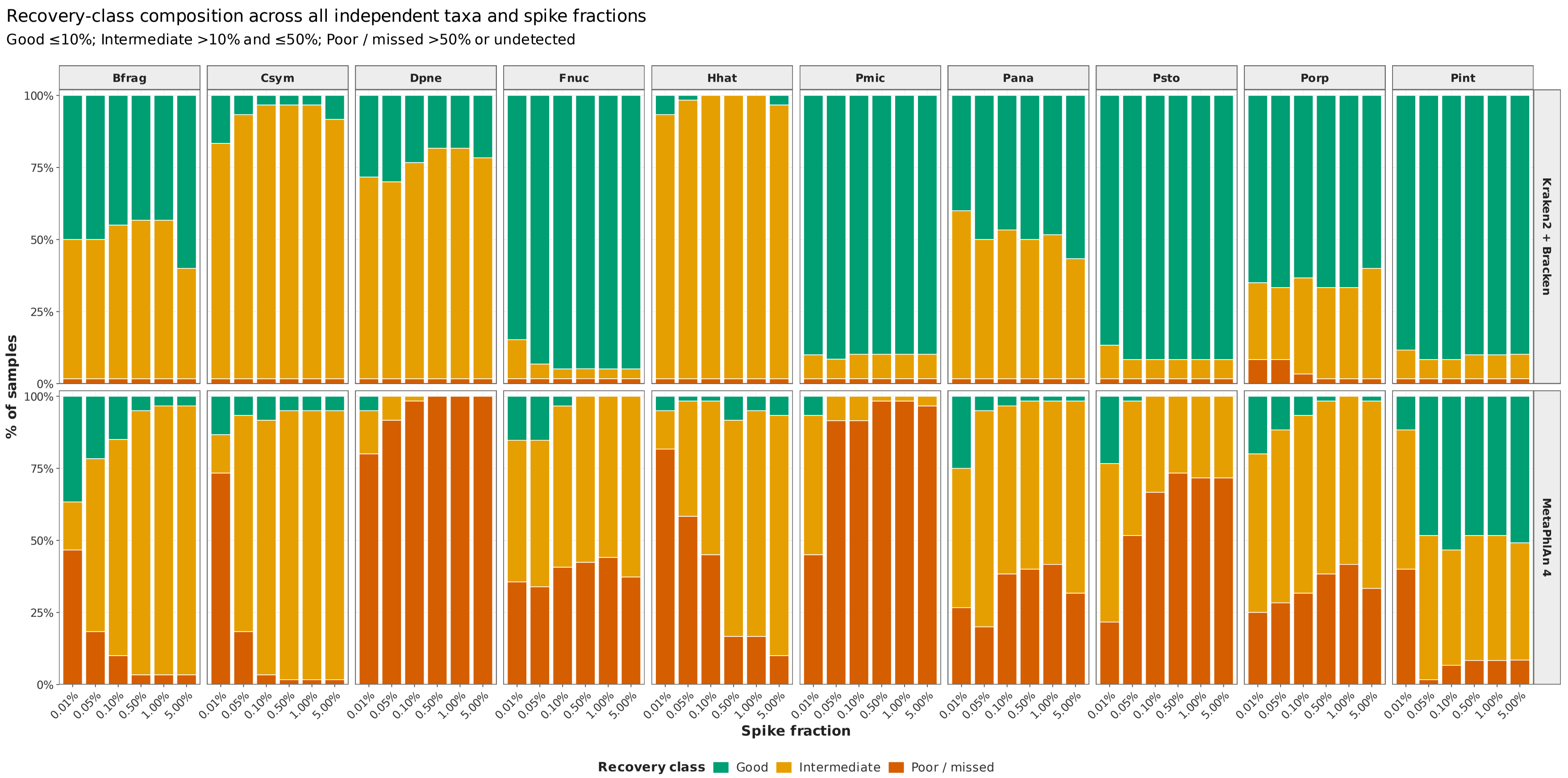


**Fig. B4: Recovery-class composition of independently spiked target taxa across all tested spike fractions.** Stacked bars show the percentage of samples assigned to each quantitative-recovery class for every target taxon, profiler, and spike fraction. Good recovery denotes an absolute relative recovery error $\leq 10\%$; Intermediate recovery denotes an error $> 10\%$ and $\leq 50\%$; and Poor/missed recovery denotes an error $> 50\%$ or failure to detect the implanted target. All ten target taxa are displayed in the manuscript-wide taxon order.

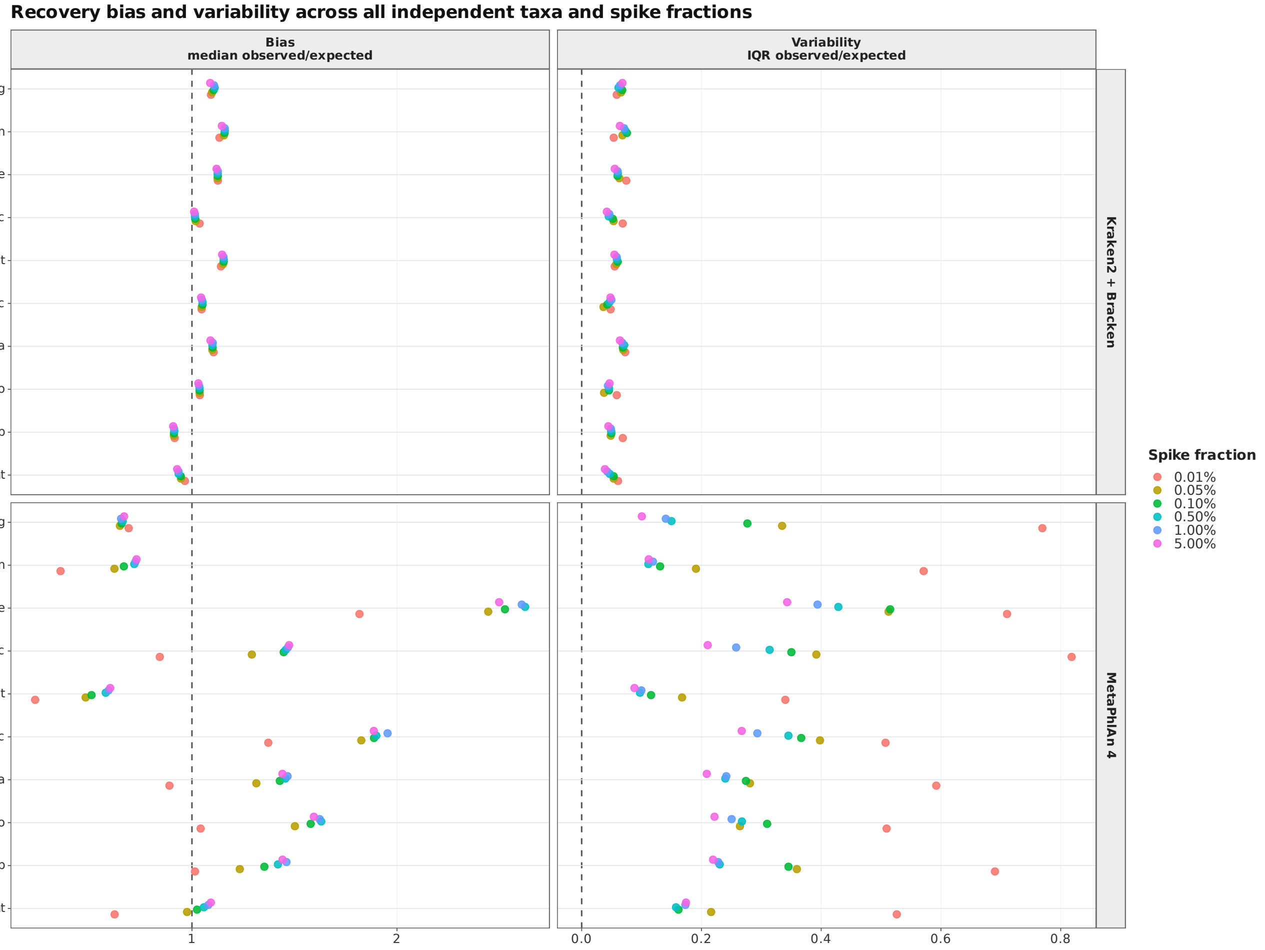


**Fig. B5: Species-level quantitative-recovery bias and variability across all independent spike fractions.** Recovery performance is summarized for each target taxon, profiler, and spike fraction. **Bias** is represented by the median observed-to-expected abundance ratio; the dashed vertical line at 1 indicates unbiased recovery. **Variability** is represented by the interquartile range of the observed-to-expected abundance ratio; the dashed vertical line at 0 indicates no between-sample variability. Colours identify the nominal spike fraction.

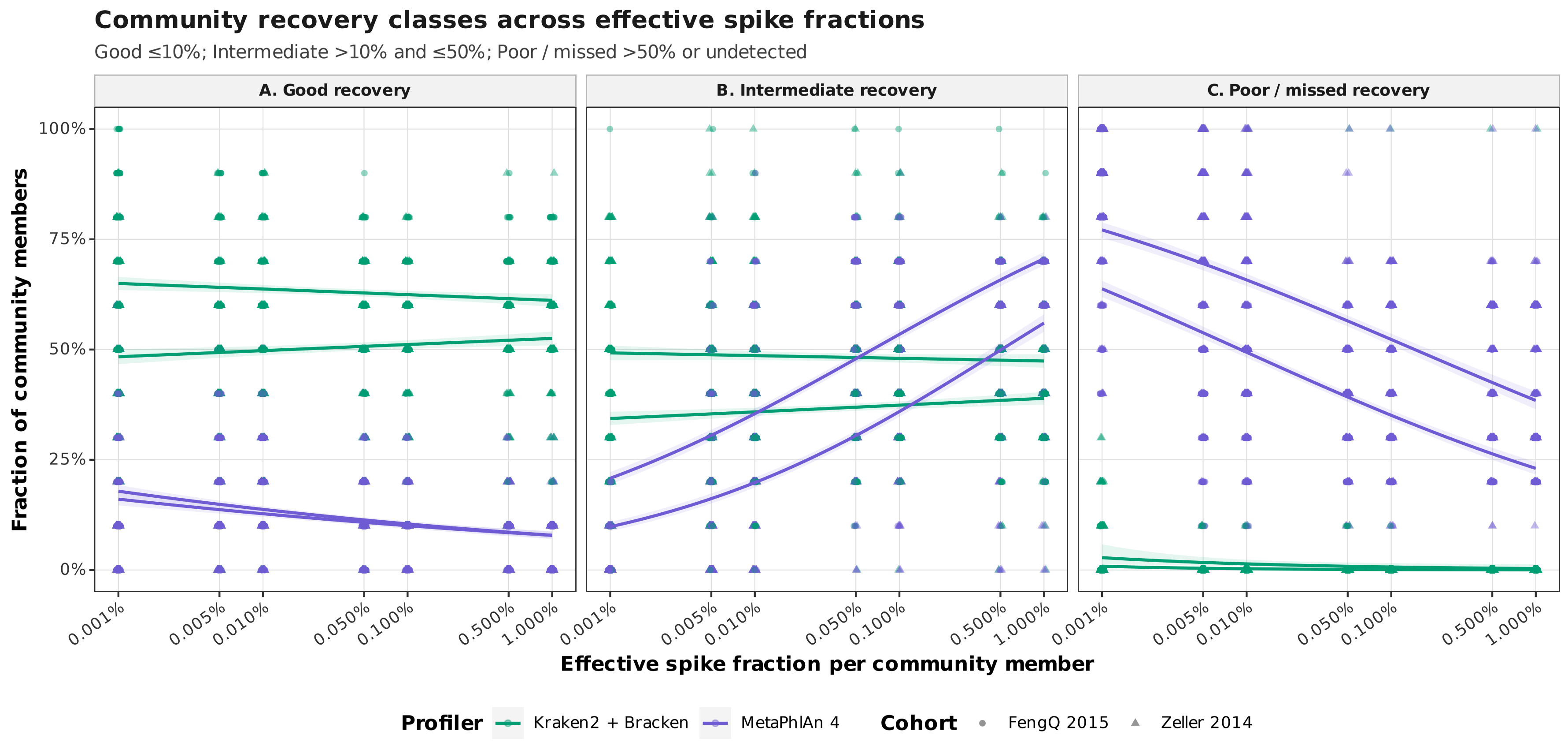


**Fig. B6: Sample-level recovery-class composition of community spike-ins across effective per-member fractions.** For each biological sample, points show the fraction of the ten implanted community members classified as **(A)** Good, **(B)** Intermediate, or **(C)** Poor/missed. Good recovery denotes an absolute relative recovery error $\leq 10\%$; Intermediate recovery denotes an error $> 10\%$ and $\leq 50\%$; and Poor/missed recovery denotes an error $> 50\%$ or failure to detect the target. Colours identify profiler and point shapes identify cohort. Lines show profiler-specific fitted trends across effective per-member spike fractions, with shaded 95% confidence intervals.

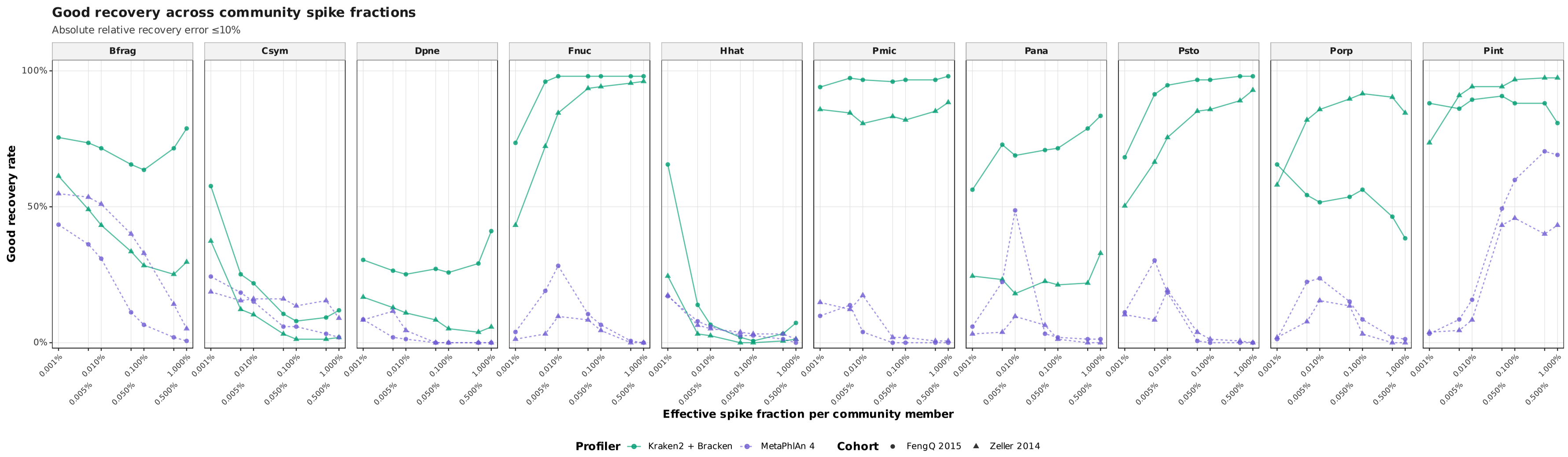


**Fig. B7: Taxon-specific Good recovery rates in community spike-ins.** Points show the percentage of samples with Good quantitative recovery for each implanted taxon across all effective per-member spike fractions. Good recovery was defined as an absolute relative recovery error $\leq 10\%$. Colours and line types identify profiler, point shapes identify cohort, and lines connect fraction-specific recovery rates within the same profiler and cohort. All ten target taxa are displayed in the manuscript-wide taxon order.

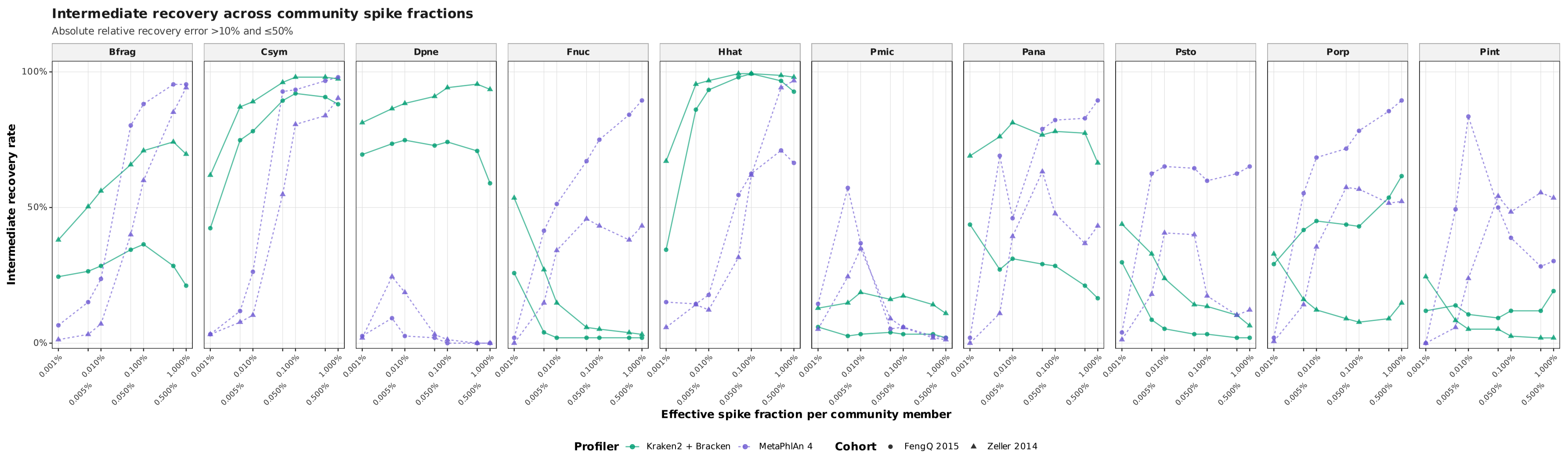


**Fig. B8: Taxon-specific Intermediate recovery rates in community spike-ins.** Points show the percentage of samples with Intermediate quantitative recovery for each implanted taxon across all effective per-member spike fractions. Intermediate recovery was defined as an absolute relative recovery error $> 10\%$ and $\leq 50\%$. Colours and line types identify profiler, point shapes identify cohort, and lines connect fraction-specific recovery rates within the same profiler and cohort. All ten target taxa are displayed in the manuscript-wide taxon order.

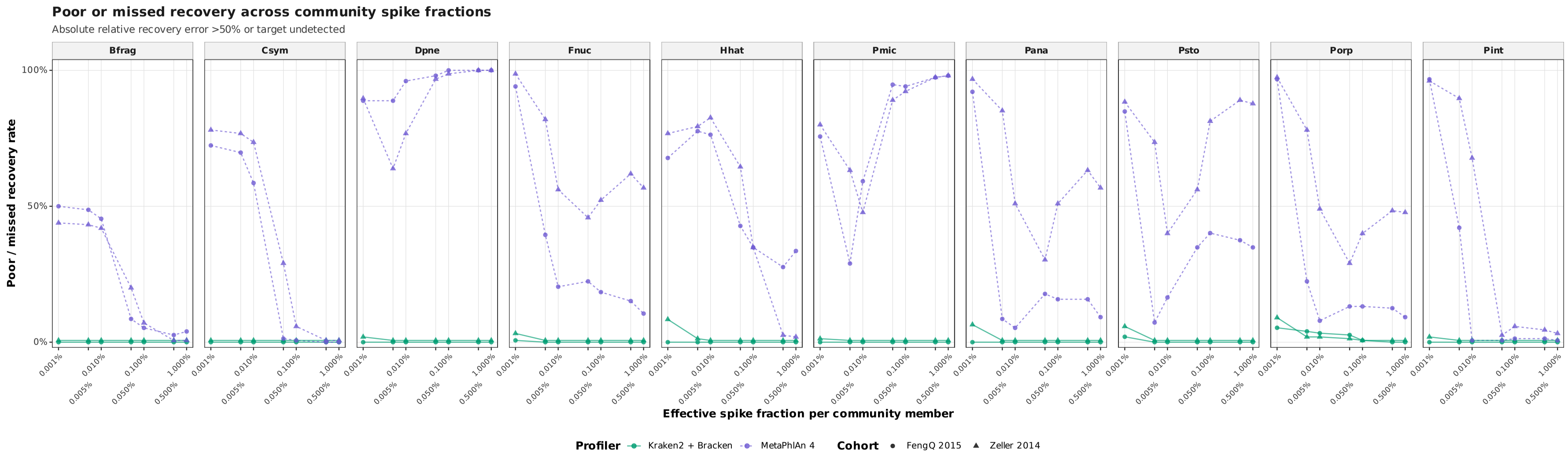


**Fig. B9: Taxon-specific Poor/missed recovery rates in community spike-ins.** Points show the percentage of samples with Poor/missed quantitative recovery for each implanted taxon across all effective per-member spike fractions. Poor/missed recovery was defined as an absolute relative recovery error > 50% or failure to detect the implanted target. Colours and line types identify profiler, point shapes identify cohort, and lines connect fraction-specific recovery rates within the same profiler and cohort. All ten target taxa are displayed in the manuscript-wide taxon order.

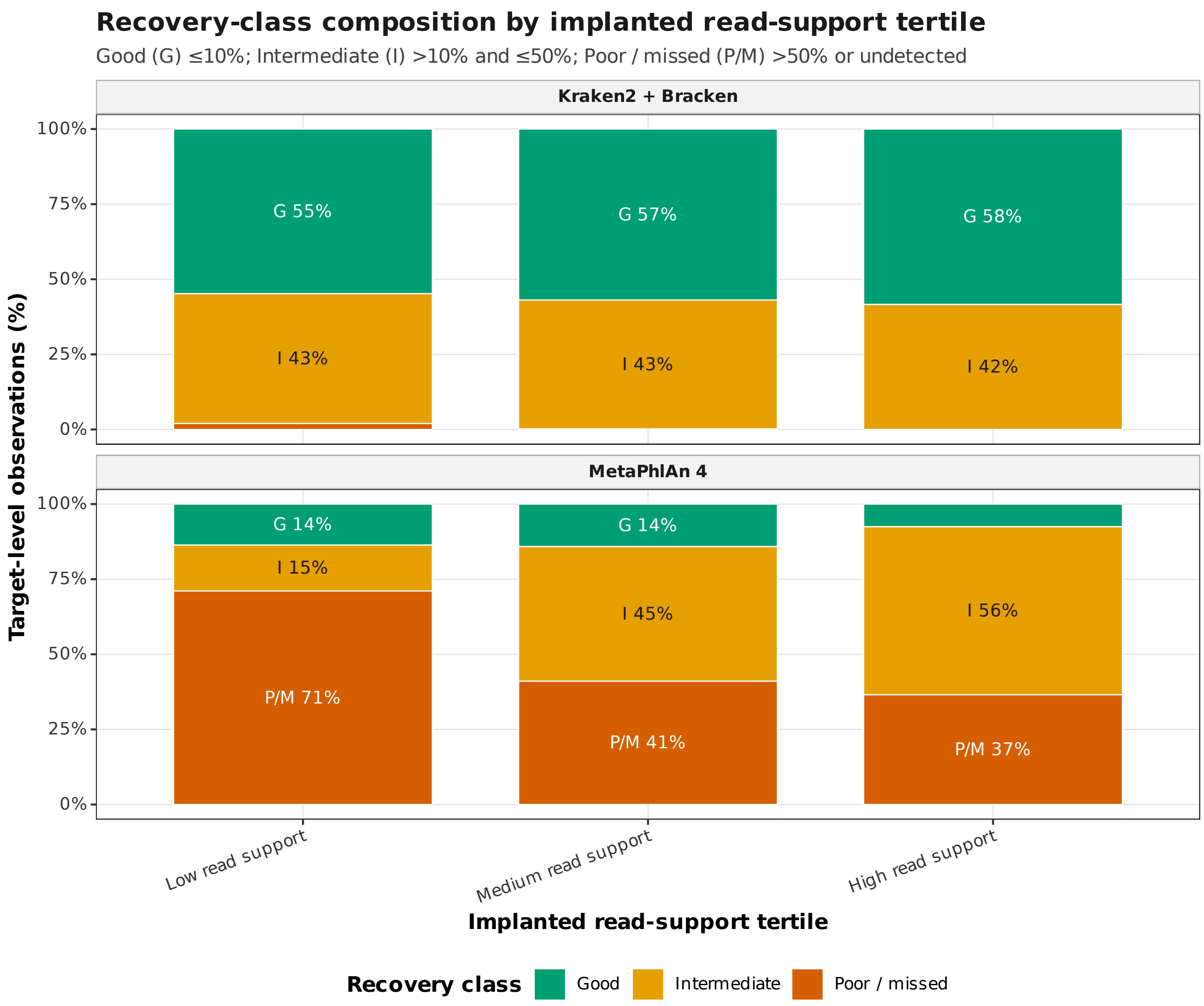


**Fig. B10: Recovery-class composition according to implanted read support.** Stacked bars show the distribution of target-level observations among recovery classes after stratification into low, medium, and high implanted read-support tertiles. Tertiles were determined from the number of implanted read pairs per target. Results are shown separately for Kraken2/Bracken and MetaPhlAn 4 and pool community-spike fractions, target taxa, cohorts, and diagnostic backgrounds within each stratum. Good (G) denotes an absolute relative recovery error $\leq 10\%$; Intermediate (I) denotes an error $> 10\%$ and $\leq 50\%$; and Poor/missed (P/M) denotes an error $> 50\%$ or an undetected target. Percentage labels identify the contribution of sufficiently large class segments.

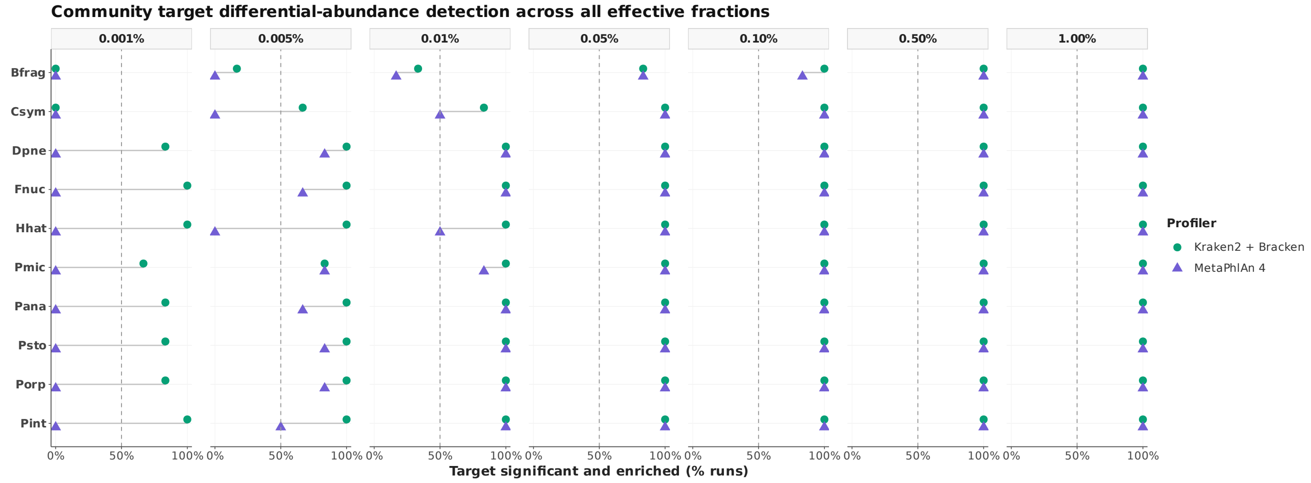


**Fig. B11: Community-spike target differential-abundance detection across all effective per-member fractions.** Points show the percentage of differential-abundance analysis runs in which each implanted target was detected as significantly enriched, separately for Kraken2/Bracken and MetaPhlAn 4. A successful target call required a positive spike-status coefficient and a false-discovery-rate-adjusted $q$-value $\leq 0.10$. Grey segments connect profiler-specific estimates for the same target and spike fraction, and the dashed vertical line marks 50% detection. Profiler estimates are slightly displaced vertically to expose overlapping values. Effective per-member fractions were obtained by dividing the total community spike fraction equally among the ten implanted taxa.

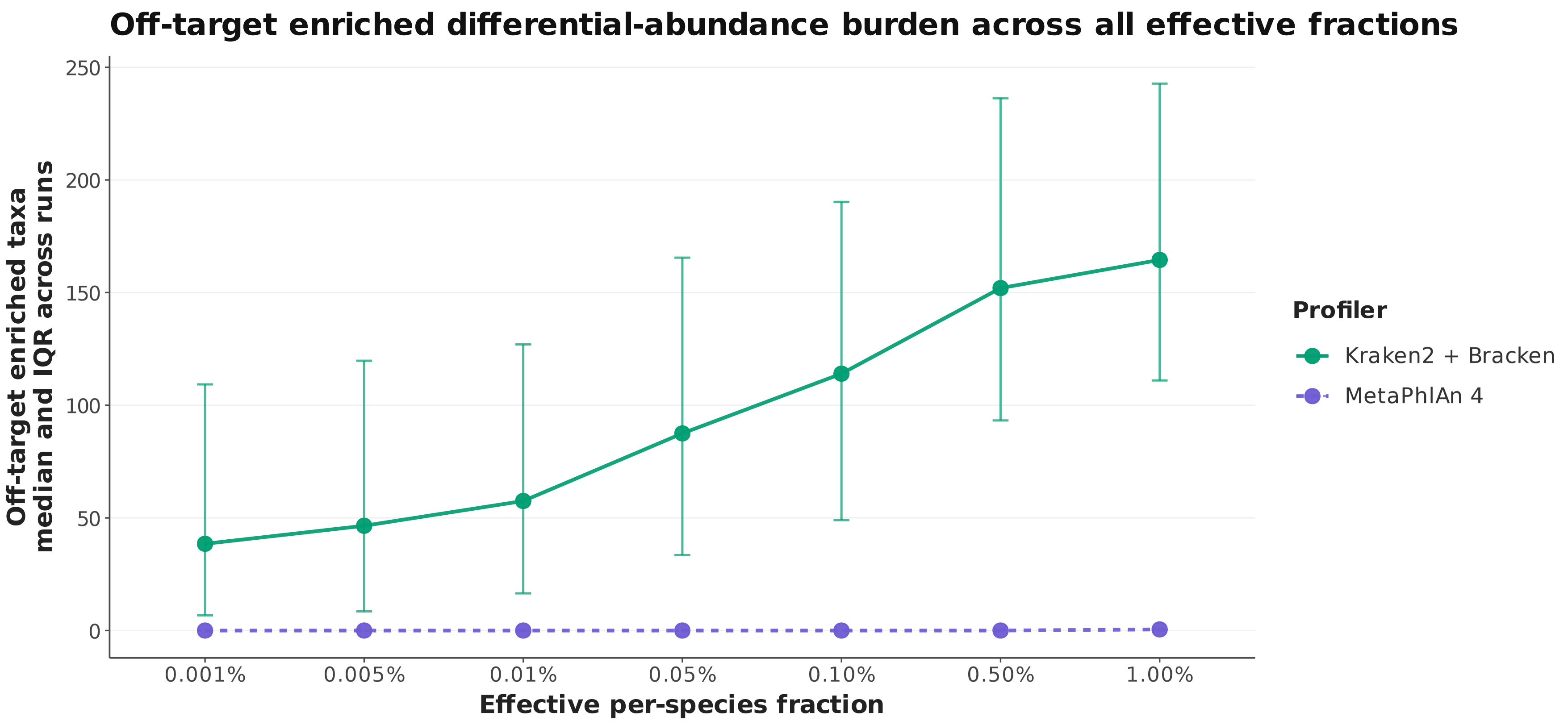


**Fig. B12: Off-target differential-abundance burden in the community-spike analysis.** Points and connecting lines show the median number of enriched non-target differential-abundance calls across analysis runs for each profiler and effective per-member spike fraction. Error bars represent the interquartile range. Enriched off-target calls were defined as non-implanted taxa with a positive spike-status coefficient and a false-discovery-rate-adjusted $q$-value $\leq 0.10$.

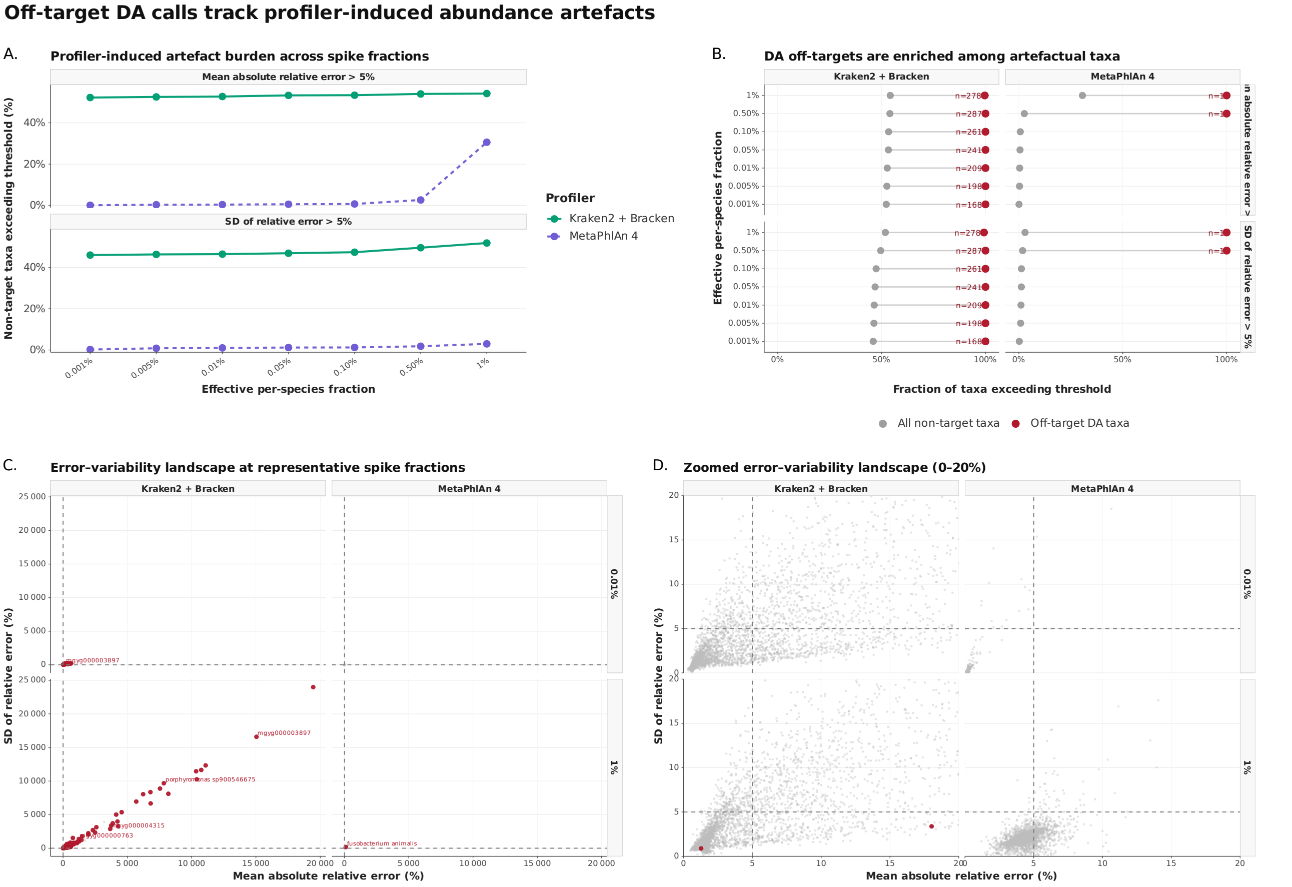


**Fig. B13: Off-target differential-abundance calls track profiler-induced abundance artefacts. (A)** Fraction of non-target taxa exhibiting profiler-induced abundance distortion across effective per-member spike fractions. Distortion is summarized using the fraction of taxa with a mean absolute relative error greater than 5% and the fraction with a standard deviation of relative error greater than 5%, calculated relative to the dilution-only expectation. **(B)** Comparison of threshold-exceedance rates among all non-target taxa and among taxa identified as enriched off-target differential-abundance calls. Grey points represent all non-target taxa, red points represent enriched off-target taxa, and connecting segments compare the two groups within each profiler, spike fraction, and error metric. Red labels indicate the number of enriched off-target taxa contributing to each estimate. **(C)** Mean absolute relative error versus the standard deviation of relative error for non-target taxa at representative effective spike fractions. Grey points represent all non-target taxa, red points identify enriched off-target differential-abundance taxa, and selected high-error taxa are labelled. **(D)** Magnified view of the error–variability landscape over the 0–20% range on both axes. Dashed lines mark the 5% mean-error and variability thresholds. Together, these analyses show that enriched off-target differential-abundance calls preferentially occur among taxa displaying profiler-induced abundance distortion.

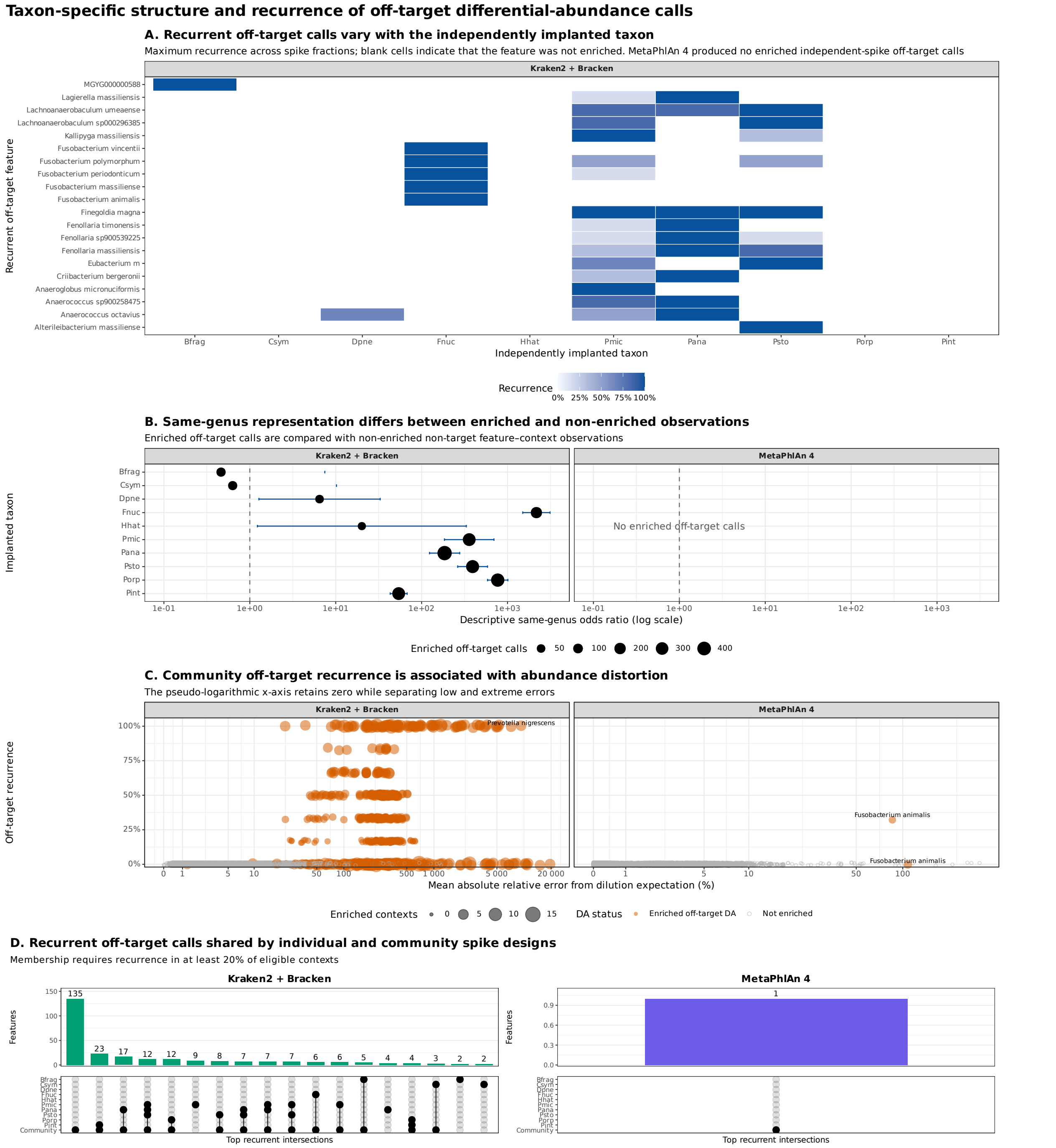


**Fig. B14: Taxon-specific structure and recurrence of off-target differential-abundance calls. (A)** Maximum recurrence of enriched off-target differential-abundance calls across individual-species spike fractions, stratified by independently implanted taxon. Recurrence denotes the fraction of eligible cohort and diagnostic-background contexts in which a non-target feature was significantly enriched. Blank cells indicate that the feature was not enriched. MetaPhlAn 4 produced no enriched individual-species off-target calls at the applied significance threshold. **(B)** Descriptive same-genus odds ratios comparing enriched off-target feature–context observations with non-enriched non-target observations. Points show odds-ratio estimates, horizontal intervals show approximate 95% confidence intervals, and point size represents the number of enriched off-target calls. The dashed vertical line denotes an odds ratio of one. Comparisons were performed separately by profiler. **(C)** Relationship between community-spike off-target recurrence and mean absolute relative error from the dilution-only abundance expectation. Point size represents the number of enriched contexts. Orange points denote features identified as enriched off-target differential-abundance features, whereas grey points denote non-enriched features. The pseudo-logarithmic horizontal scale retains zero while separating low and extreme errors. **(D)** Intersections of recurrent off-target features between the ten individual-species spike-in experiments and the community-spike experiment, shown separately by profiler. Membership required recurrence in at least 20% of eligible contexts. Bars show the number of features belonging to each intersection, and the matrix identifies the contributing spike-in designs.
